# Design and Construction of Three Customizable Pilot-Scale Hydroponic Systems

**DOI:** 10.64898/2026.09.24.747602

**Authors:** Liam Reynolds, Brandon Hollenback, Paul Davidson

## Abstract

An increasing number of researchers, educators, and at-home growers are turning to hydroponics as an alternative method of food production, diversifying and strengthening the food supply chain. This has led to a substantial increase in the amount of academic work involving hydroponic production of plants for a variety of purposes, including but not limited to a more controlled and customizable environment than conventional production methods, utilizing hydroponics as a method of wastewater treatment, and investigating the vertical production of plants on a much smaller footprint than traditional commercial-scale production. Because so many unique stakeholders are interested in hydroponics for a variety of reasons, academic studies use a wide variety of hydroponic systems with varying levels of technology and automation, making comparisons across studies and systems challenging. A few institutions have sophisticated plant growth and research facilities, like those used by commercial companies, but these systems are often prohibitively expensive, costing upwards of $10,000 USD for a very limited amount of growing space for experimental trials. To enable more researchers, educators, and citizen scientists to conduct experiments and easily compare their results, this study details three pilot-scale hydroponic production systems (Kratky, NFT, and DWC), utilizing entirely off-the-shelf components, for under $5,000 USD each, that are capable of maintaining the desired consistent temperature and relative humidity levels, and have space for 216, 108 (6 irrigation loops), and 64 (16 irrigation bins) plants, respectively. The material lists and prices are provided along with assembly instructions and general guidelines for each hydroponic system. The testing and optimization process of environmental conditions within each growth system is also detailed, with suggested customizations for potential experimental setups. The authors developed these systems during their Master’s and Doctoral degree programs, enabling improved hydroponic research at the University of Illinois Urbana-Champaign. They hope that other aspiring hydroponic researchers can benefit from this work in the development and operation of their hydroponic systems, increasing research efficiency and collaboration opportunities in this field of study.

## 1. Introduction & Objectives

There is an emerging need to adapt current food production practices to overcome the challenges of climate change (Mirzabaev et al., 2023), population growth, and urbanization (Ackerman, 2012). The agricultural industry is facing a growing challenge of producing more food (Van Dijk et al., 2021) with fewer inputs (Grafton et al., 2015) on less land (Rudel, 2021) while ensuring there is still enough potable water for society to survive (Karpe et al., 2025).

Urban agriculture is uniquely positioned to produce food near or directly in urban environments where most consumers are located, reducing food miles and the time from farm to table (Nowysz et al., 2022). Controlled environment agriculture takes this even further with the capability to produce food year-round within greenhouses or vertical farms (Vatistas et al., 2022). The latter of which also leverages what would otherwise be unused vertical space to produce more food per unit area than traditional agriculture (I. Abdurakhmonov, 2025).

Urban and controlled environment agriculture are not without their own challenges (Kalantari et al., 2018), including higher capital and operating costs due to the necessary infrastructure and electrical power consumption (Specht et al., 2014), in addition to competing with higher urban land values (Orsini et al., 2013). Private companies and academic institutions have dedicated R&D funds and personnel to improve the CEA industry (list examples of each to give them their dues), and there has been progress in recent years, but there is still a barrier to entry because of the high upfront capital costs and the complexity in keeping all the various components and considerations of a CEA system up and running (Shamshiri, Kalantari, et al., 2018). These challenges can prevent smaller companies or academic institutions from entering the CEA space (Burritt et al., 2025), slowing down the pace of R&D and, therefore, slowing the growth and improvement of the industry. To combat these entry challenges, cost-effective and easy-to-operate lab-scale hydroponic systems must be developed.

The ideal conditions for hydroponic crop growth and development vary across plant species and the type of growth system utilized, just as they do for non-hydroponic crop production (Namany et al., 2025). Therefore, any pilot-scale hydroponic system needs to maintain a wider range of environmental conditions to accommodate a wide range of individual crop species. The system is then adjusted to the specific production conditions necessary for that particular crop at that stage of its growth and development. The critical factors affecting plant growth and development in CEA, besides the availability of nutrients, are light intensity and duration (light integral), the amount of moisture in the air (relative humidity or vapor pressure deficit), and the temperature of the air and leaf (Boros et al., 2023; Shamshiri, Jones, et al., 2018). CEA is a system of systems, and as such, it will always require fine-tuning and optimization to the specific conditions and contexts of the grower.

CEA systems generally allow for the adjustment of light intensity and duration, which together form the light integral, circulation fan rate and placement, exhaust fan rate and placement, and humidifier rate and placement. These systems work in concert to create the ideal microclimate around the plant canopy and root zone, as these factors drastically affect plant growth and development (Santosh & Shukla, 2024). Circulation fans are utilized to mix the air immediately around the plant leaves, enabling better plant respiration and ensuring minimal spatial environmental differences (Zhang & Kacira, 2018). Many of these factors work against each other in a production setting because of the additional heat generated by artificial lighting and the humidity load of crop respiration. For example, when the lights are on and generating heat, it can become too hot for ideal plant growth, necessitating exhaust fans to turn on or increase their rate to expel this hotter air and intake cooler, external air. However, this cooler external air is often less humid than necessary for plant growth, so the humidifiers must turn on or increase their output to accommodate this additional humidification load.

There is a wide range of laboratory or greenhouse-scale hydroponic systems in the scientific literature. These can consist of simple jars filled with a nutrient solution, atop which a Styrofoam media floats with a plastic wrap lid (Aslam et al., 2023), to miniaturized commercial production systems (Nikolov et al., 2023) and state-of-the-art plant growth chambers used in specialized facilities. The wide range of growth systems means just as wide a range of growth conditions and operations are needed, making it difficult to make comparisons between and across studies. Cost-effective and representative hydroponic research systems need to be standardized in terms of the environmental conditions they can maintain as well as the degree of control an operator has over those conditions. Additionally, the specific research interests of the Circular Bioeconomy Resource group in the ABE department at UIUC, focused on the reuse of alternative water sources as nutrient sources for hydroponic crop production, require growth systems that can accommodate a few different irrigation waters being applied, compared to most existing growth systems at this scale that only have one reservoir source water.

Therefore, the specific objectives of this study are to:

1. Create pilot-scale hydroponic growth systems that provide control over light intensity, humidity, wind speed, and temperature.
2. Tabulate the components and costs of these systems, enabling other researchers to replicate them; and
3. Detail the tuning process to adjust the conditions to the desired ranges for specific experimental needs.

## 2. Methods

Three different hydroponic growth systems were designed and constructed to test the effects of alternative irrigation waters on hydroponic crop production: 1. A microgreen production system that doubles as a germination system, 2. a DWC grow tent, and 3. an NFT grow tent. These three systems are specifically designed for testing many different irrigation waters in a small footprint, enabling research on every stage of a crop cycle, from germination and initial growth (microgreens) to vegetative growth (leafy greens & herbs) to fruiting and flowering (edible flowers and fruiting crops).

The microgreen system was developed as a preliminary growth chamber capable of supporting germination of crops to be used in the other two systems, as well as to provide a simple, controlled environment capable of supporting many experimental replicates. The compact growing levels enable screening experiments to investigate many different crops and irrigation solutions within the same overall growth chamber. The DWC system was developed to explore plants that require a larger area for growth or have more intensive root systems than the other systems could accommodate. Flowering species such as sunflowers have demonstrated remediation characteristics, which would be appropriate for alternative irrigation source water exploration, but also require a large area for shoot growth and housing for the large taproot produced. A large reservoir provided by a DWC system would be suitable for these types of applications. The NFT system was developed to align with many commercial hydroponic systems, as the thin nutrient film enables excellent oxygenation of the root zone, while maintaining required moisture levels. This system is suitable for leafy greens and herbs that do not have an expansive root system, as the compact growing levels and irrigation trough volumes cannot support flowering or fruiting species.

Many of the same pieces of equipment are used across all three systems, including a 971 Temperature Humidity Meter from Fluke (Everett, WA, USA) for handheld spot measurements, multiple Drop D2 Wireless Temperature and Humidity Data Loggers from Kestrel (Albuquerque, NM, USA) for continuous data collection, a LED LT40 light meter from Extech (Nashua, NH, USA) for preliminary lux measurements, an LI-180 Spectrometer from LI-COR (Lincoln, NE, USA) for direct PPFD and red:blue:green:far-red light measurements, a 405i Hot Wire Anemometer from Testo (Titisee-Neustadt, Germany) for wind speed measurements, and finally the LV600HH Hybrid Ultrasonic Humidifier from Levoit (Anaheim, CA, USA) are used to provide humidity in all three growth systems.

A common lighting schedule in CEA operations is 16-hour days with 8-hour nights, and the minimum PPFD should reach at least 200 µmol/m2/s, giving an ideal DLI of approximately 12 mol/m2/day (Yan et al., 2019). The typical ideal temperature range for hydroponics is between 18-27 °C, while the ideal relative humidity is between 50-70% (Naranjani et al., 2022). The ideal wind speed to provide enough air exchange for photosynthesis is greater than 0.2 m/s (Kitaya et al., 2004).

### Germination/Microgreen Growth Chamber

The microgreen and germination growth systems consist of three 1.22 m W x 1.83 m H x 0.61 m D metal shelving racks, creating nine growth levels, each with a volume of approximately 283.2 L available for plant growth, while maintaining sufficient airflow. These metal shelving racks are enclosed with a reflective wrapping, the Cool Shield Thermal Bubble Roll from Uline (Pleasant Prairie, WI, USA), to create a thermal and moisture barrier, allowing the space inside the chambers to be better controlled and semi-isolated from the ambient lab conditions. This wrapping does not create a perfect seal, as flaps were cut for the ventilation fans along the top of the chambers, along with flaps cut along the back wall of the chambers to access electrical outlets. Additionally, a series of corrugated black plastic sheets make up the floor of the lowest growth levels, allowing the humidified air to remain within the plant environment more effectively. The reflective wrapping, covering every side of the metal shelving racks, is secured mostly with duct tape; however, simple binder clips are used to hold the front flaps in place, creating a growth chamber with 9 growth levels, while allowing easy access to the plants for data collection and observations. In Figure 1 below, the empty microgreen system is pictured with the middle flap rolled open, and the system is unpowered.

**Figure 1.**
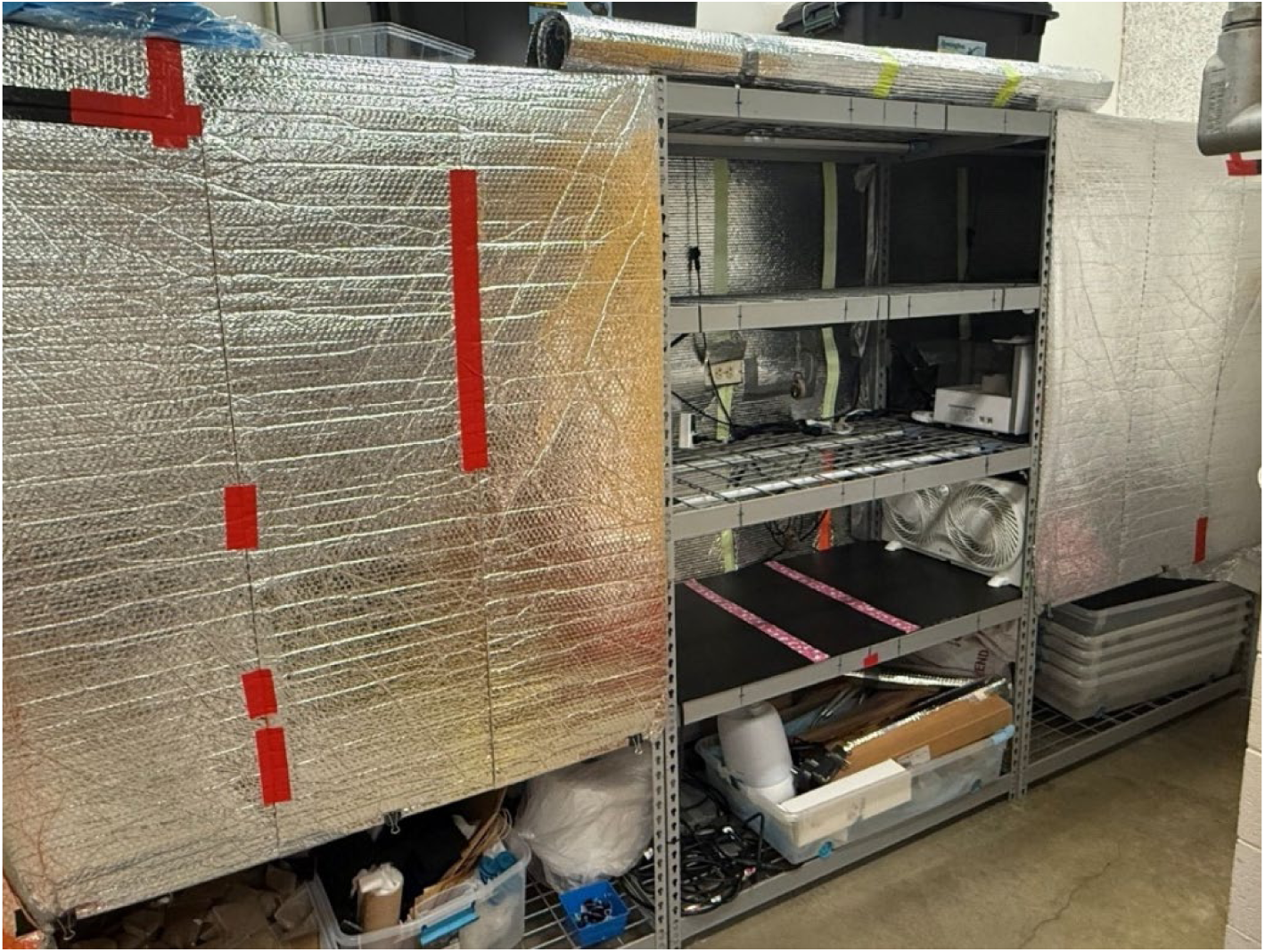
Image of empty hydroponic microgreens and germination growth chamber with the middle flap raised.

A humidifier and circulation fan are positioned between each of the three metal racks, where they meet, with the humidifier above the circulation fan to prevent accidental water accumulation within the fans if the humidifier expels air directly into the fans’ intake side. The circulation fans, 22.86 cm window fans from Utilitech (Jacksonville, NC, USA), are arranged where the direction of each half of the fan, the fan allows for the left and right sides to be controlled independently, alternates with the other to better mix the air within the growth racks, and the direction of the humidifier output splitters alternates as well, ensuring the same environmental conditions are experienced on each of the 9 growth levels. There is one ventilation fan, a 12.7 cm square USB computer fan from AC Infinity (City of Industry, CA, USA) in the center of the ceiling of each of the three metal growing racks, and temporary flaps are used to cover these fans to prevent excess air exchange during the dark germination stage, as the ventilation fans are only needed to pull the excess heat generated by the LED lights away from the plant environment. There are two Strata light bars on each of the 9 growth levels, located 26 cm apart, 12.1 cm from the front/back edges. A technical drawing of the microgreen growth chamber is shown below in Figure 2, to scale. A picture of the system in use is provided (Figure 9) as an additional reference, and a table detailing the quantity and cost of components for the construction of the system is provided (Table 1).

**Figure 2.**
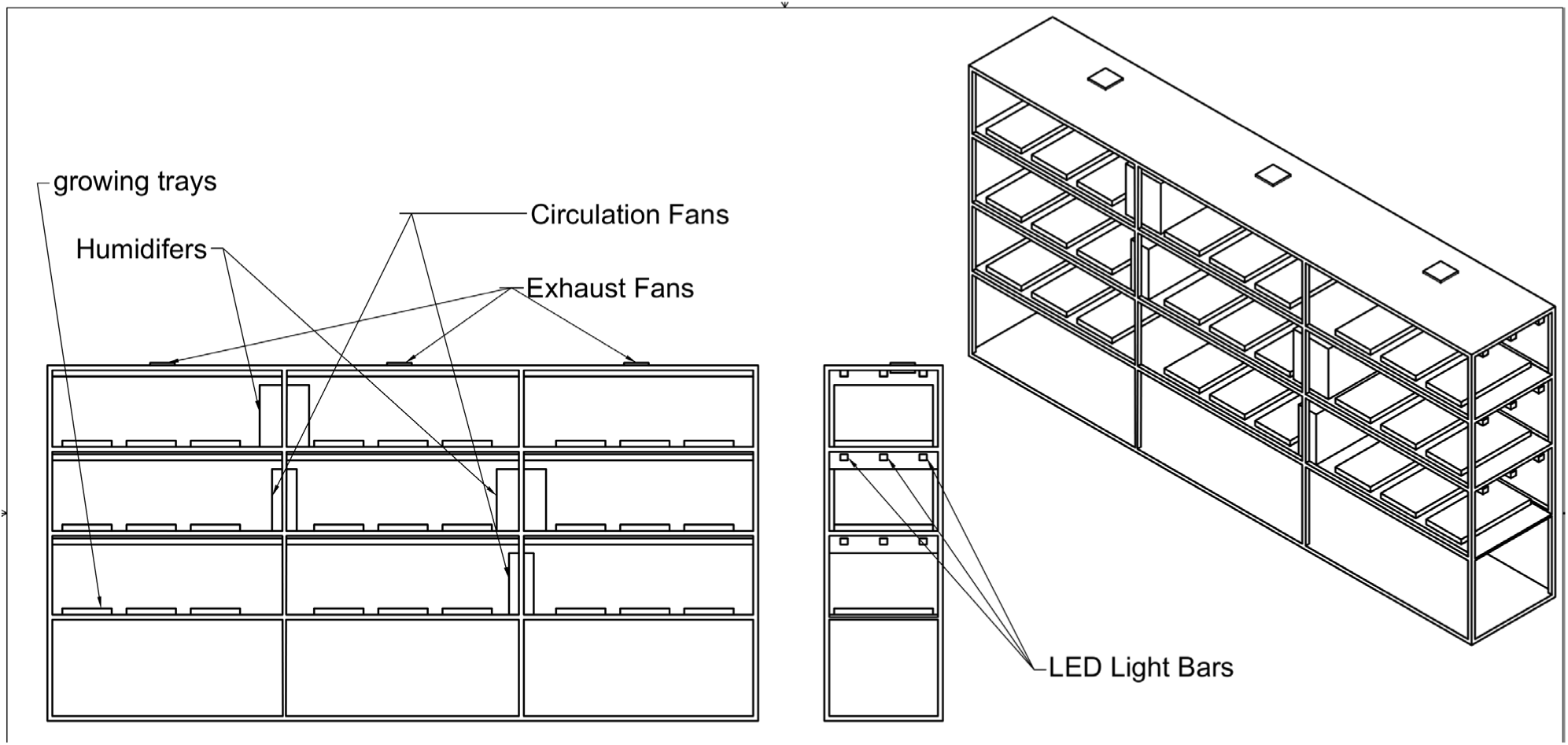
Technical drawing of hydroponic microgreens and germination growth system to scale.

**Table 1.**
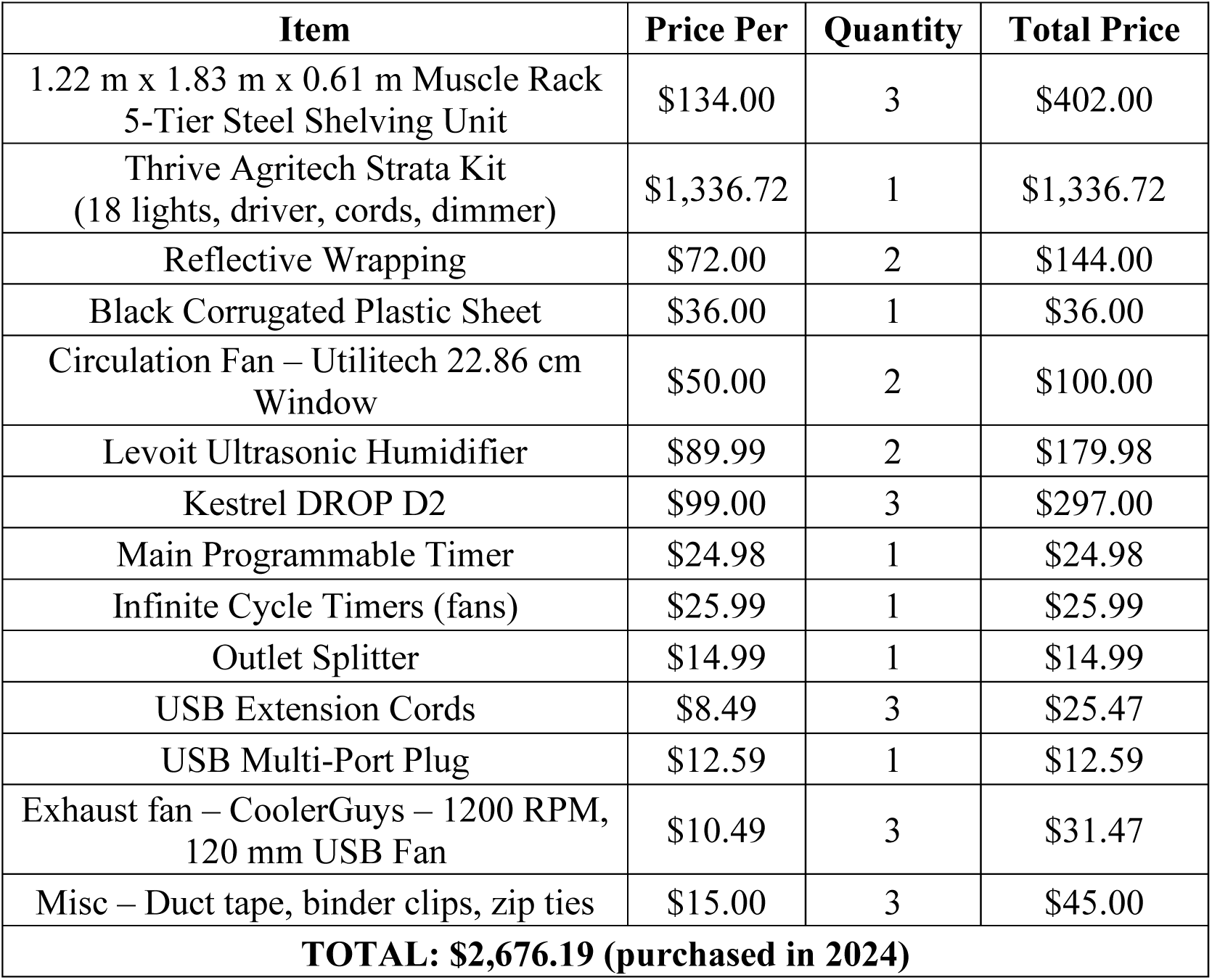
Materials and costs to replicate microgreen and germination growth system.

### Deep Water Culture Growth Chamber

The Deep-Water Culture Growth System consists of a 1.98 m W x 1.98 m H x 1.98 m D grow tent from Grow Ace (Anaheim, CA) with three metal shelving units at 1.22 m W x 1.83 m H x 0.61 m D from Menards (Eau Claire, WI), each with the capacity of five levels, but only two in use on each for optimal full-growth height spacing. The lighting consists of 12 Thrive Agritech (New York, NY) Infinity XE models at 1.14 m W x 0.08 m H x 0.06 m D, with 168 μmols/sec light output at a 120-degree beam angle. These lights operate at a 16:8 photo period controlled by the timer provided by Thrive Agritech. The top and bottom levels have multiple strips of Cool Shield Thermal Bubble Roll from Uline (Pleasant Prairie, WI, USA) attached to improve lighting reflectivity and ensure uniformity. The strips of reflective sheeting each measure 0.42 m W x 1.59 m H. The large height allows the full strip to span the top and bottom levels. Placement of the sheeting is indicated in Figure 3 below by dashed lines.

**Figure 3.**
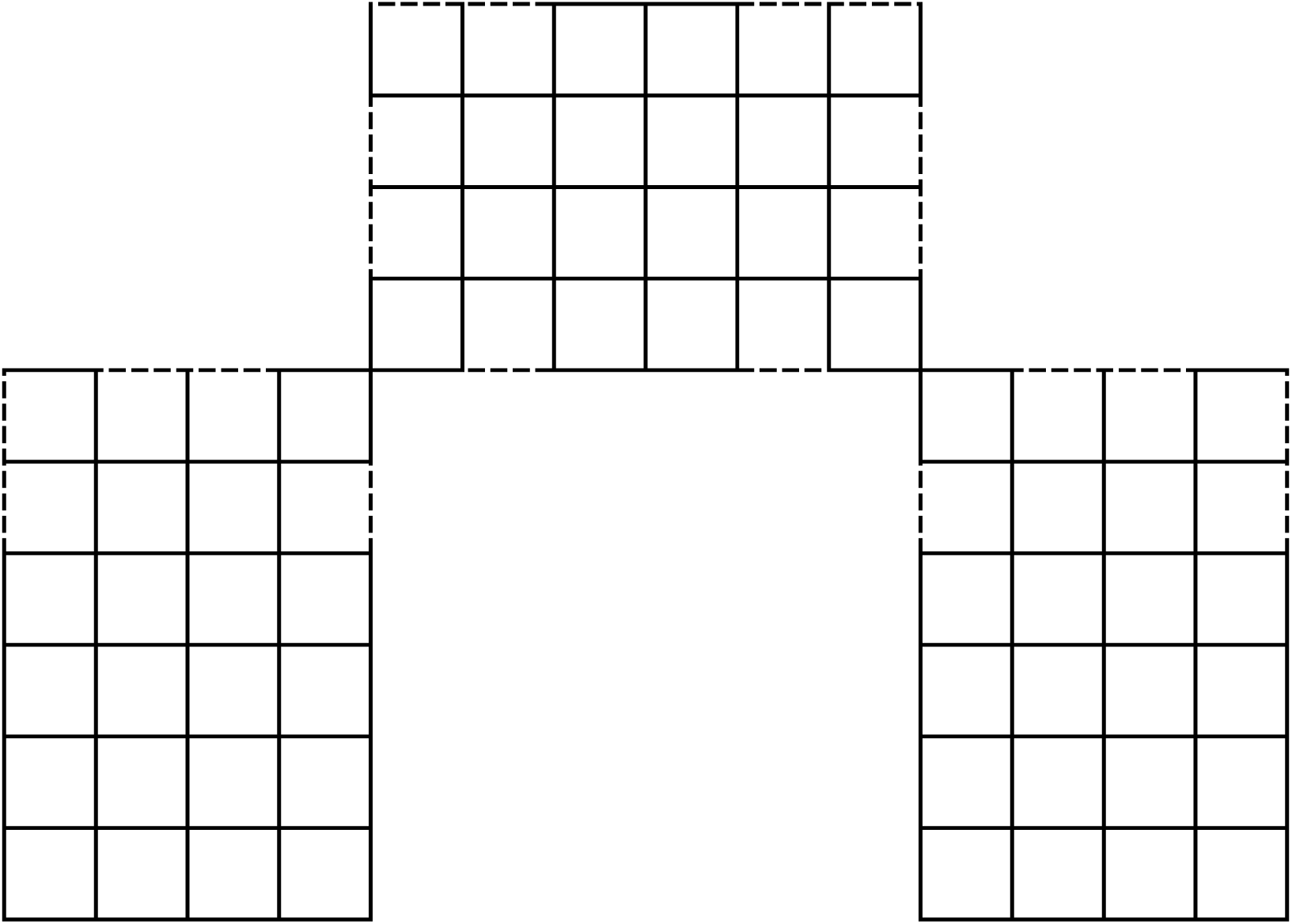
Dashed lines indicate where the reflective sheeting is installed on both the top and bottom levels.

Each level accommodates approximately 17 cubic feet for growth (0.61 m W x 0.65 m H x 1.22 m D). Germination and seedling growth are performed with Gardzen 40-cell seed trays (City of Industry, CA). Each seed tray measures 0.37 m W x 0.05 m H x 0.23 m D, with the humidity dome adding 0.06 m in height. Up to 8 plug trays can fit comfortably on a single level, with five placed vertically and three placed horizontally. When the full DWC system is in place, three Rubbermaid (Atlanta, GA) Roughneck 10-gallon totes fit on each level, totaling 15 possible unique irrigation solutions, each with a capacity for four plants via holes drilled in the lids. Each tote uses a large cylindrical air stone with air provided from a pump with a 45 L/min output, split with a six-outlet manifold, to provide approximately 7.5 L/min of aeration to the plant roots. Each line of tubing (5 mm inner diameter, 7 mm outer diameter) is cut to the length needed to reach the middle inside the storage tote.

For airflow, dual 120 mm fans from AC Infinity (City of Industry, CA) are installed on each level, in alternating directions, to provide optimal air circulation and uniformity, with 2.95 m^3^/min per set. The total air movement with these fans is 17.67 m^3^/min. Additional air circulation is provided via a Pelonis 1.02 m oscillating tower fan from Menards (Eau Claire, WI). For the exhaust, a VIVOSUN R6 0.15 m Inline Fan (Ontario, CA, USA) is adhered to the roof on the outside of the tent, with a pull rate of 11.04 m^3^/min. The fan is connected to two vents located in the back corners of the tent via a T-splitter, enabling approximately 5.52 cubic meters of air to be removed from each vent in the growth system and released into the ambient environment. No active intake is present; only four ventilation holes are opened on the front left and right sides, allowing for slight negative pressure. Proper airflow also ensures uniformity with humidity, which is supplied by four Levoit (Anaheim, CA) Ultrasonic 6L LV600HH-RXW humidifiers, each set at a 75% humidity level. The temperature and humidity levels are verified to be within optimal ranges by utilizing six Kestrel Instruments (Boothwyn, PA) Drop D2 data loggers, one positioned hanging in the center of each growing level. All electronics are stored in a 10-gallon clear storage tote with a lid to ensure protection from water. Figures 4 and 5 below detail each growth rack to scale. The cost of development is provided in Table 2 below.

**Figure 4.**
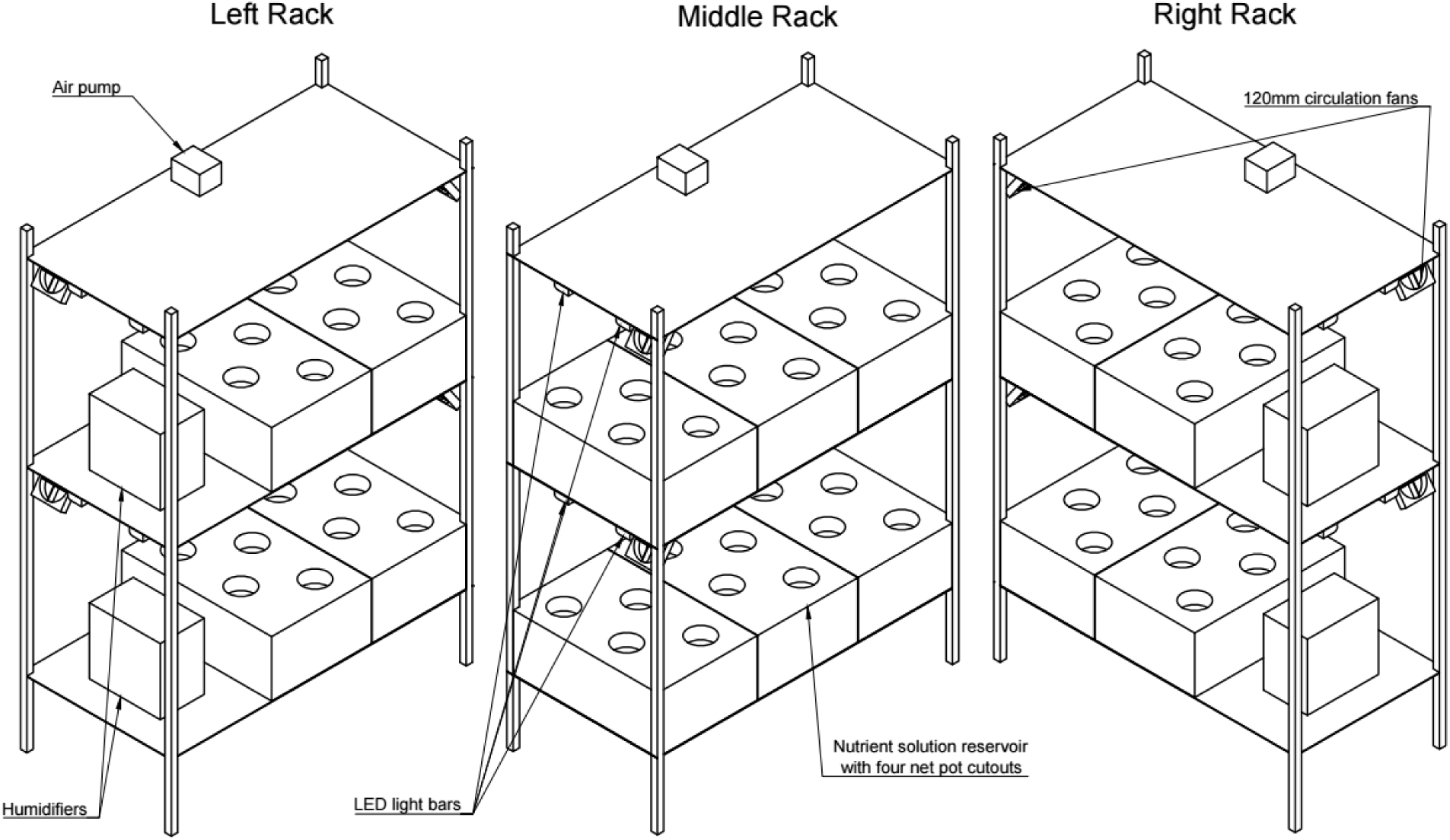
Technical drawing of the DWC growth racks to scale in different arrangements.

**Figure 5.**
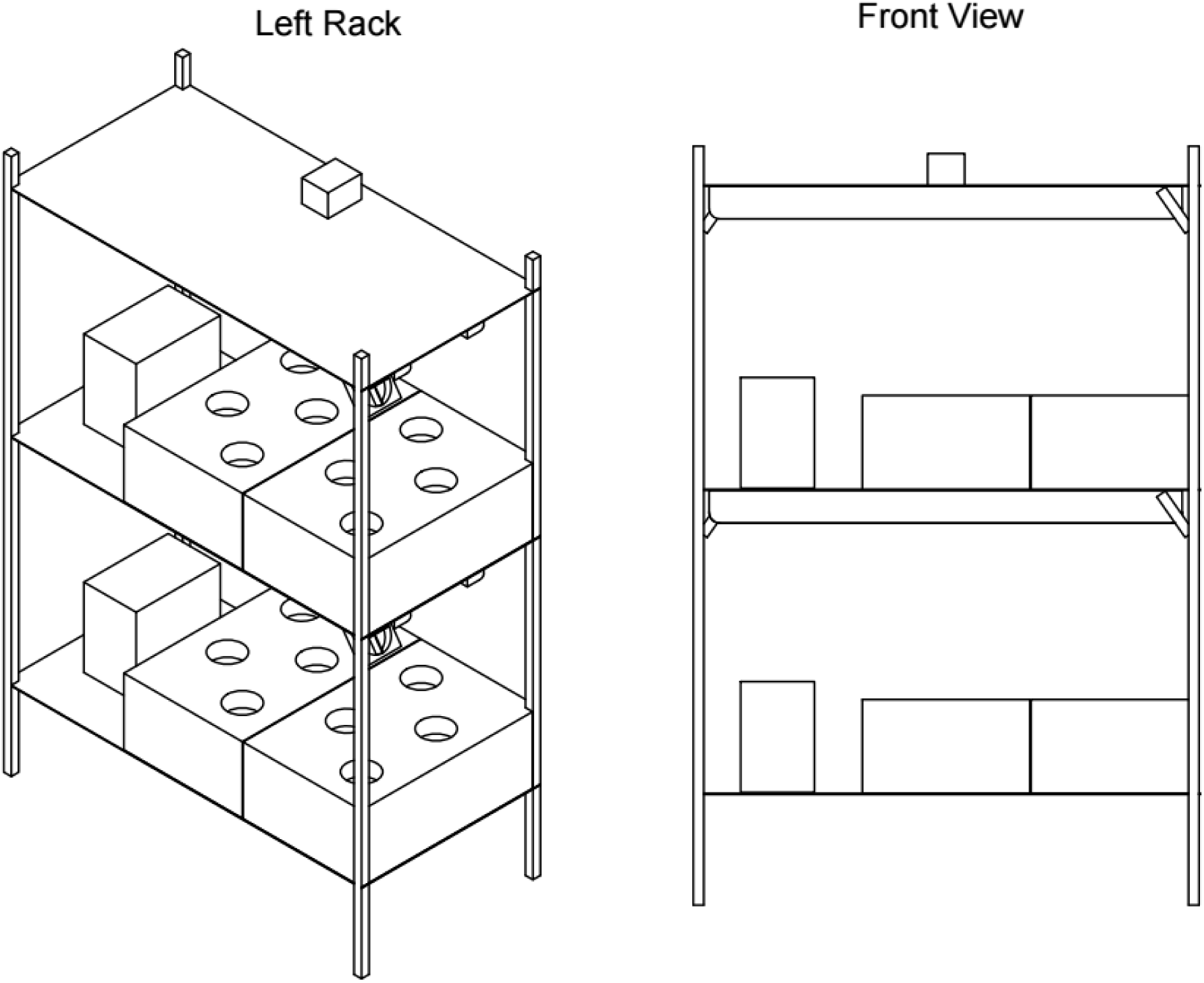
Technical drawing of the DWC growth chamber’s left rack with an additional side view, to scale.

**Table 2.** Materials and costs of the DWC growth chamber.

| **Item** | **Price Per** | **Quantity** | **Total Price** |
| --- | --- | --- | --- |
| Yield Lab Reflective Grow Tent –<br>1.98 m x 1.98 m x 1.98 m | \$202.47 | 1 | \$202.47 |
| Muscle Rack 1.22 m W x 1.83 m H x<br>0.61 m D<br>5-Level Steel Shelving Unit | \$134.00 | 3 | \$402.00 |
| Thrive Agritech Infinity XE Lighting<br>Kit (lights, driver, cords,<br>timer/dimmer) | \$150.00 | 12 | \$1,800.00 |
| Reflective Wrapping | \$72.00 | 2 | \$144.00 |
| Rubbermaid Roughneck Tote 10 Gal<br>- 6 Pack | \$80.99 | 3 | \$242.97 |
| Levoit Ultrasonic Humidifier – 6L | \$89.99 | 3 | \$269.97 |
| Kestrel DROP D2 | \$99.00 | 3 | \$297.00 |
| Vivosun - R6 Inline Fan 0.15 m with<br>Variable Speed Controller | \$79.99 | 1 | \$79.99 |
| Vivosun – 0.15 m x 2.44 m Black<br>Non-Insulated Flex Air Aluminum<br>Ducting | \$15.99 | 1 | \$15.99 |
| Interior fan – AC Infinity -<br>MULTIFAN S7 - Dual 120mm | \$19.99 | 6 | \$119.94 |
| Pelonis 1.02 m Oscillating Tower Fan | \$49.99 | 1 | \$49.99 |
| Extension Cables | \$9.98 | 3 | \$29.94 |
| Surge Protector | \$9.88 | 1 | \$9.88 |
| Miscellaneous – duct tape, zip ties,<br>binder clips | \$15 | 1 | \$15 |
| **TOTAL: \$3,679.14 (purchased in 2024)** | | | |

### Nutrient Film Technique Growth Chamber

The recirculating NFT Growth System consists of a 1.83 m W x 2.44 m H x 1.22 m D grow tent from Grow Lights Canada (Lincoln, ON, Canada) that holds two 1.22 m W x 1.83 m H x 0.61 m D metal shelving racks, creating six growth levels, each with a volume of approximately 283 L available for plant growth, and each capable of testing a different irrigation solution. The grow tent creates a thermal and moisture barrier, allowing the space inside to be isolated from the ambient environment. However, it is not a perfect seal, as there are very minor gaps around the zippers and the ducting ports in the tent; for temperature control, two of the four intake ventilation flaps are open. The ventilation exhaust pulls from both the top left and right sides of the growth chamber, with the 15.2 cm ducting centered above the top level of each of the growth racks. The ventilation unit is a 15.2 cm 11,043.6 L/min inline fan from Vivosun (Ontario, CA, USA) that is connected to a T-splitter to pull approximately 5,521.8 L of air from each side before exhausting into the ambient space. This ventilation creates a small amount of negative pressure, another reason for opening two of the four bottom ventilation flaps. There are two 12.7 cm square USB computer fans from AC Infinity (City of Industry, CA, USA) on each of the 6 growth levels to provide sufficient air circulation between the NFT growth troughs on each level. As in the other growth systems, the circulation fans are arranged in alternating directions for better air mixing and uniformity of environmental conditions. There is one humidifier placed in the middle of the chamber, furthest from the entrance, between the two growth racks. There are three Strata light bars on each of the 6 growth levels, located 16.5 cm apart, 10.2 cm from the front/back edges. Corrugated black plastic serves as the base of each growth level to prevent light from another level from affecting results, in addition to providing a flatter surface for the NFT troughs to be uniformly inclined on.

The 10.2 cm wide, 2.44 m long NFT troughs were purchased from GrowSpan (South Windsor, CN, USA) and cut in half to create 18 1.22 m long troughs, capable of supporting 5 plants each, for 15 plants per growth level, totaling up to 90 plants that can be grown in the tent at one time. The 6^th^ plant slot is too close to the outlet, and roots may clog it, so it was not used; additionally, around 2.54 cm of water pools around the outlet, another reason the 6^th^ plant slot was not used. Growspan recommended a 3 – 4 cm rise over the 1.22 m trough run to ensure sufficient water movement. To achieve this slope, trough risers were 3-D printed to raise the inlet side by 3.175 cm. Additionally, spacers were 3-D printed to ensure the troughs stayed straight and aligned with the grow lights above them. These spacers are 7.62 cm long, and the.stl files of the risers and spacers are provided in the Appendix. Each growth level is irrigated from an 18.9- liter food-safe bucket located below the growth levels, and an adjustable 33.3 L/min pump from Hydrofarm (Pennsylvania, USA) is used for each of the two highest levels to accommodate the extra elevation head, while a non-adjustable 25 L/min pump from EcoPlus (Charlotte, NC, USA) is used for each of the other 4 levels. The water is pumped through 2.54 cm diameter food-safe tubing until it reaches a DIY 3-way splitter created with standard food-grade PVC components. Three 0.64 cm tubes connect the splitter to the three NFT troughs on each level, where it runs the length of the troughs before flowing out into 2.54 cm food-safe tubing, into a 3-way junction, before being returned to the 18.9-liter irrigation reservoir, oxygenating the water. A technical drawing of the NFT growth chamber system is shown below in Figure 6, to scale. A picture of the system in use is provided in Figure 16 as an additional reference. Important to note that a 37.85-liter plastic tote is utilized to store all electronic components, preventing accidental spills or leaks from shorting the system and creating a safety hazard. A table detailing the quantity and cost of components for the construction of the system is provided (Table 3).

**Figure 6.**
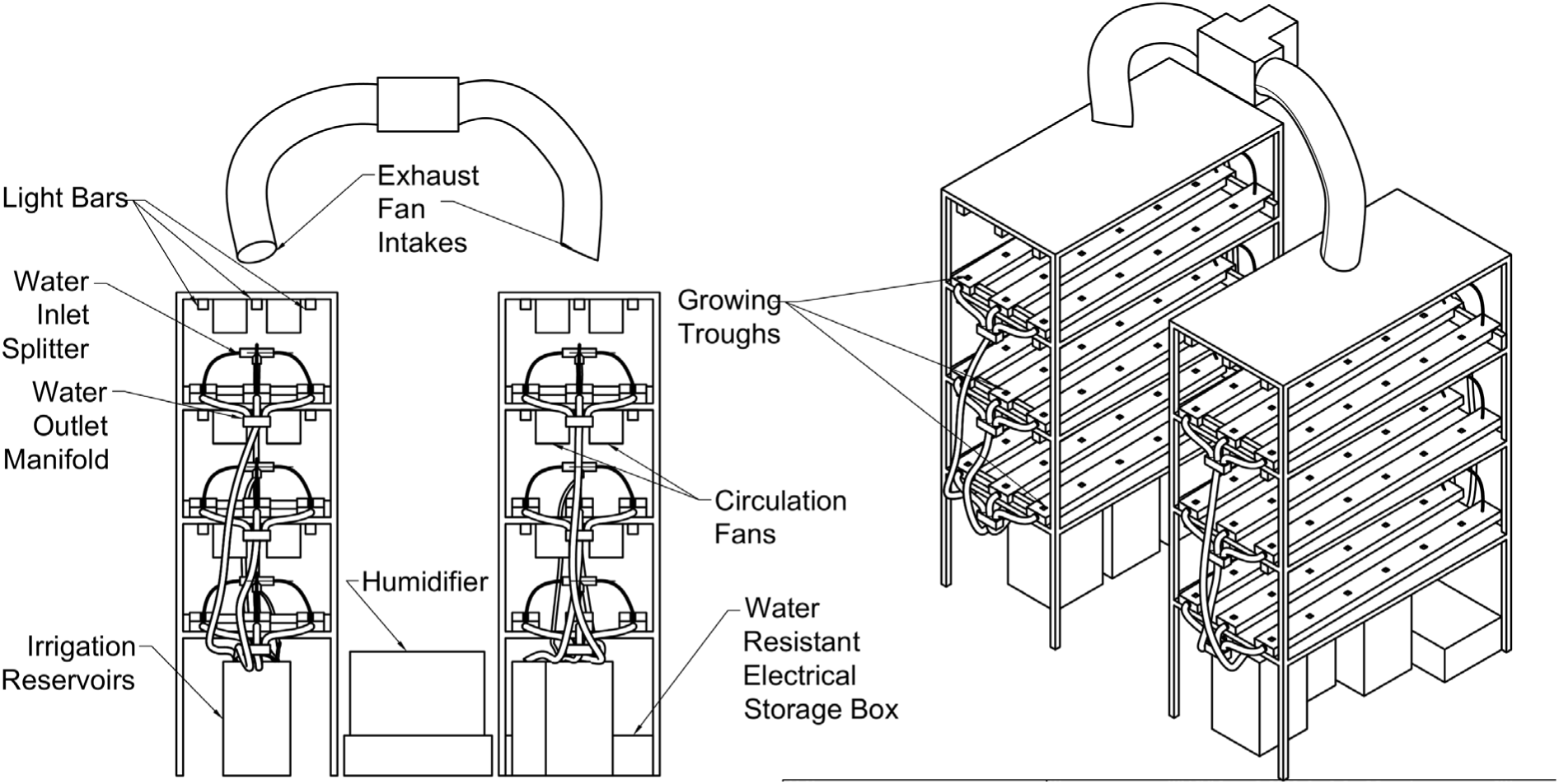
Technical drawing of the NFT hydroponic system, approximately to scale.

**Figure 6:**
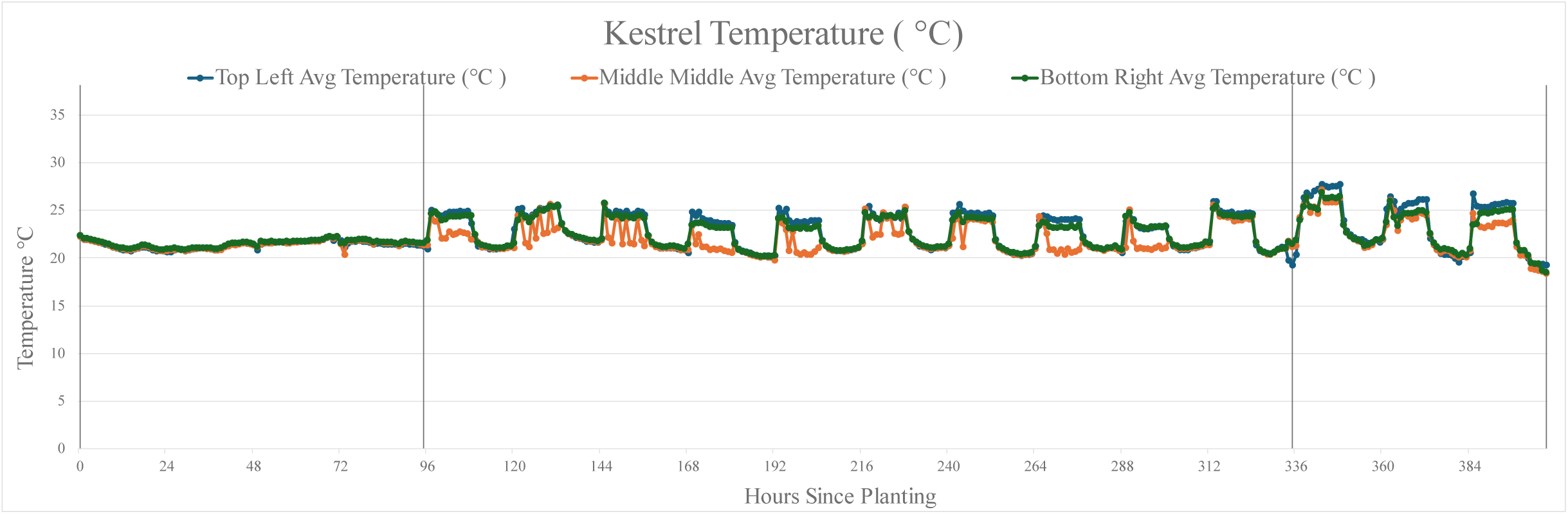
Kestrel temperature data from the 7th microgreens experiment.

**Table 3.** Materials and costs to replicate NFT growth system.

| **Item** | **Price Per** | **Quantity** | **Total Price** |
| --- | --- | --- | --- |
| Fusion Hut Reflective Grow Tent –<br>1.83 m x 2.44 m x 1.22 m | \$209.00 | 1 | \$209.00 |
| 1.22 m x 1.83 m x 0.61 m Muscle<br>Rack 5-Tier Steel Shelving Unit | \$134.00 | 2 | \$268.00 |
| Thrive Agritech Strata Kit<br>(18 lights, driver, cords, dimmer) | \$1,336.72 | 1 | \$1,336.72 |
| GrowSpan NFT Package<br>(troughs, 0.64 cm tubing, shipping) | \$600.00 | 1 | \$600.00 |
| Black Corrugated Plastic Sheet | \$36.00 | 2 | \$72.00 |
| 37.85-liter buckets and lids | \$13.90 | 6 | \$83.4 |
| 33.3 L/min submersible pump | \$49.93 | 2 | \$99.86 |
| 25 L/min submersible pump | \$35.96 | 4 | \$143.84 |
| Levoit Ultrasonic Humidifier | \$89.99 | 1 | \$89.99 |
| Kestrel DROP D2 | \$99.00 | 3 | \$297.00 |
| Main Programmable Timer | \$24.98 | 1 | \$24.98 |
| 2.54 cm diameter food-grade tubing | \$91.54 | 3 | \$274.62 |
| 4-way Electrical Outlet Splitter | \$14.99 | 2 | \$29.98 |
| USB Extension Cords | \$8.49 | 6 | \$50.94 |
| USB Multi-Port Plug | \$12.59 | 4 | \$50.36 |
| Ventilation Inline Fan - Vivosun -<br>11,043.6 L/min<br>(also ducting & clamps) | \$95.98 | 1 | \$95.98 |
| Circulation fan – CoolerGuys – 1200<br>RPM, 120 mm USB Fan | \$10.49 | 12 | \$125.88 |
| Extension Cord | \$9.98 | 3 | \$29.94 |
| Surge Protector | \$9.98 | 1 | \$9.98 |
| PVC Misc – inlet Tee, caps, outlet<br>junction, hose clamps,<br>threaded/barbed connectors | \$40.00 | 6 | \$240 |
| Misc – Duct tape, superglue, zip ties,<br>silicone | \$15.00 | 4 | \$60.00 |
| **TOTAL: \$4,192.47 (purchased in 2024)** | | | |

## 3. Results

### Microgreen/Germination Grow Chamber

The microgreen and germination chamber has been developed and optimized for our research needs over 7 microgreen growth experiments, in addition to 2 germination experiments, where the seedlings were grown in preparation for transplanting into the NFT system. The results of each experiment led to improvements, including the reflective wrapping, the installation of exhaust ventilation fans, and the optimization of the humidifier and circulation fan settings. Over the course of these experiments, the average germination and growth temperatures were reduced to acceptable levels, while efforts were made to increase the humidity levels in the growth chamber while maintaining sufficient airflow and air exchange. The Kestrel Datalogger temperature and relative humidity hourly data from the latest experiment are plotted below in Figures 6 and 7, with the germination and growth periods separated by a vertical black line. The germination temperature and relative humidity averaged 21.6 ± 0.6 °C and 72.5 ± 7.3%, while the growth temperature and relative humidity averaged 23.6 ± 0.1.7 °C and 61.5 ± 11.2%.

**Figure 7:**
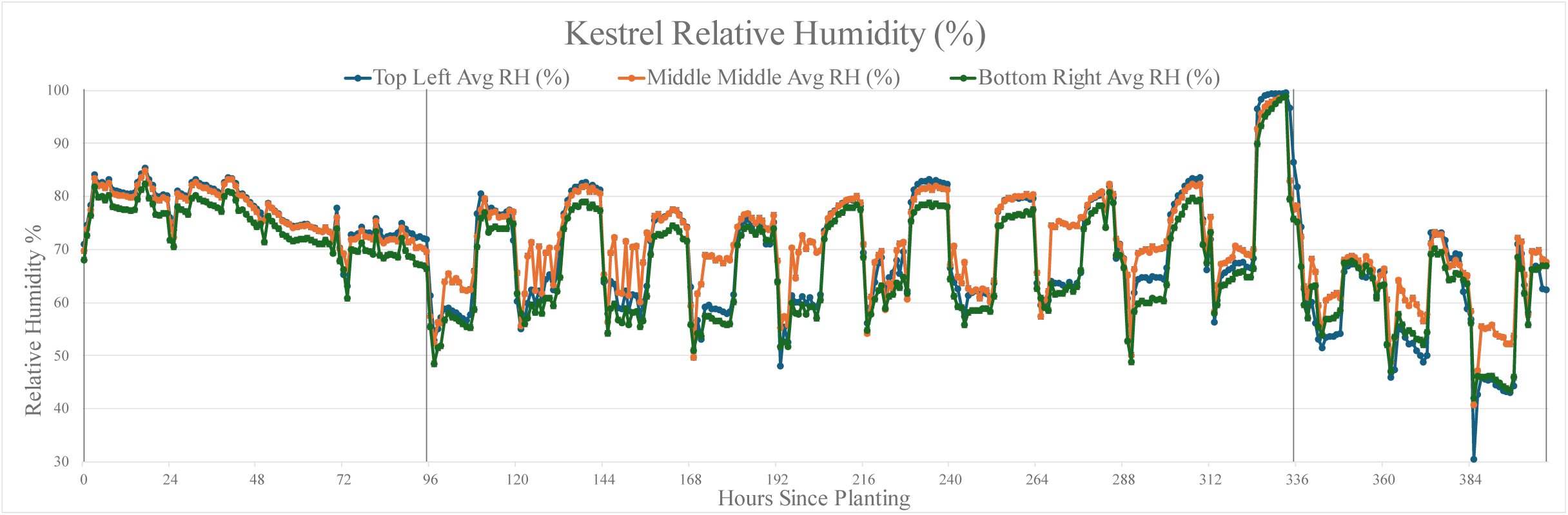
Kestrel relative humidity data from the 7th microgreen experiment.

The light intensity (in units of PPFD) was mapped in the microgreen grow chamber according to the following arrangement: 4 measurements were taken along the center of each 25.4 cm x 50.8 cm black tray, corresponding to where each microgreen replicate would be. There are three black trays per growing level, so there are 16 measurements for a given height and light setting per growth level. Measurements were taken at 50%, 75%, and 100% light intensity settings at heights of tray level, 15.24 cm above the tray, and 30.48 cm above the tray (or just below the LED light bar). This measurement arrangement ensured the light intensity (PPFD) variability is known across a given growth level and at the light intensity settings that are utilized in various experimental stages. The light intensity was also measured at various distances between the bottom of a growth level and the bottom of the LED light bars, as the plants will grow taller during the experiment, increasing the light intensity they receive. The relevant light intensity maps for microgreen production, 100% intensity at tray level and 15.24 cm above tray level, are provided in supplementary Tables 1 and 2, but the average PPFD over the whole microgreen growth chamber is 160.17 ± 8.17 for tray level, and 186.88 ± 12.02 for 15.24 cm above. These light intensity levels are appropriate for microgreen growth and seedling germination, but higher levels are recommended for full-scale crop growth, giving reason for the higher light intensity in the DWC and NFT growth tents.

The wind speed was measured once for each black 25.4 cm x 50.8 cm tray in the center of the tray at the average crop canopy height throughout the microgreens’ growth experiments. There are definite “hot spots” of higher wind speed directly near the two circulation fans and “cold spots” of lower wind speed; this is one of the main factors that influences the rotation of crops and treatments between the 9 different growth shelves, as it ensures all replicates experience the same conditions on average. The overall microgreens grow chamber average wind speed was 0.17 ± 0.21 m/s, and the wind map is shown in supplementary Table 3. This average wind speed is slightly below the minimum of 0.2 m/s necessary for uniform air mixing in some levels within the grow chamber, again highlighting how important the rotation of replicates is, especially around the crop canopy, facilitating gas exchange and evapotranspiration, enabling the photosynthesis that powers crop growth. All the environmental conditions per growth level within the microgreen growth chamber are averaged and presented in Table 4 below. Figure 8 depicts the microgreen growth chamber in use during one of the microgreen experiments, with all three front flaps open to reveal the layout of the experimental replicates.

**Figure 8.**
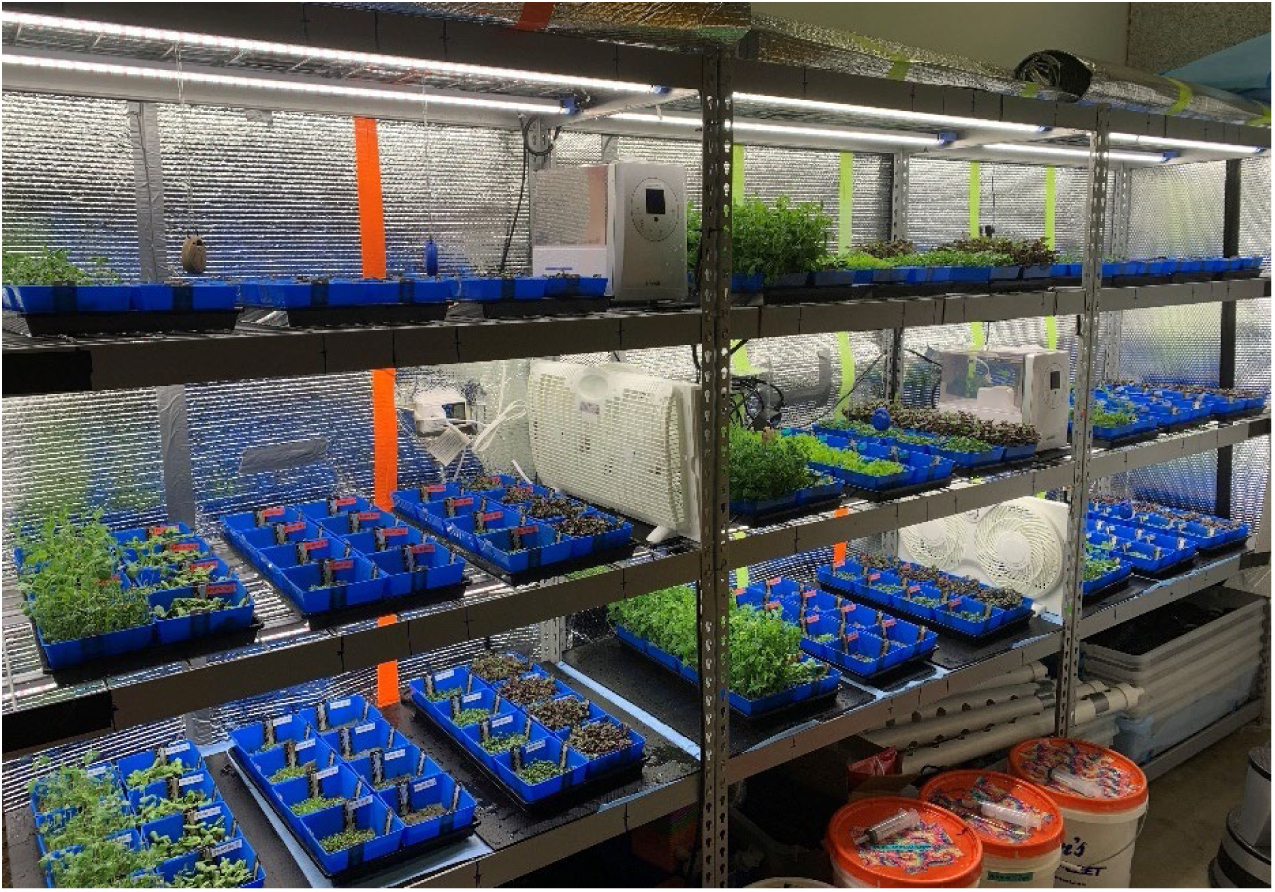
Image of the Microgreen Growth Chamber in Use with the front flaps completely open for visibility.

**Figure 9.**
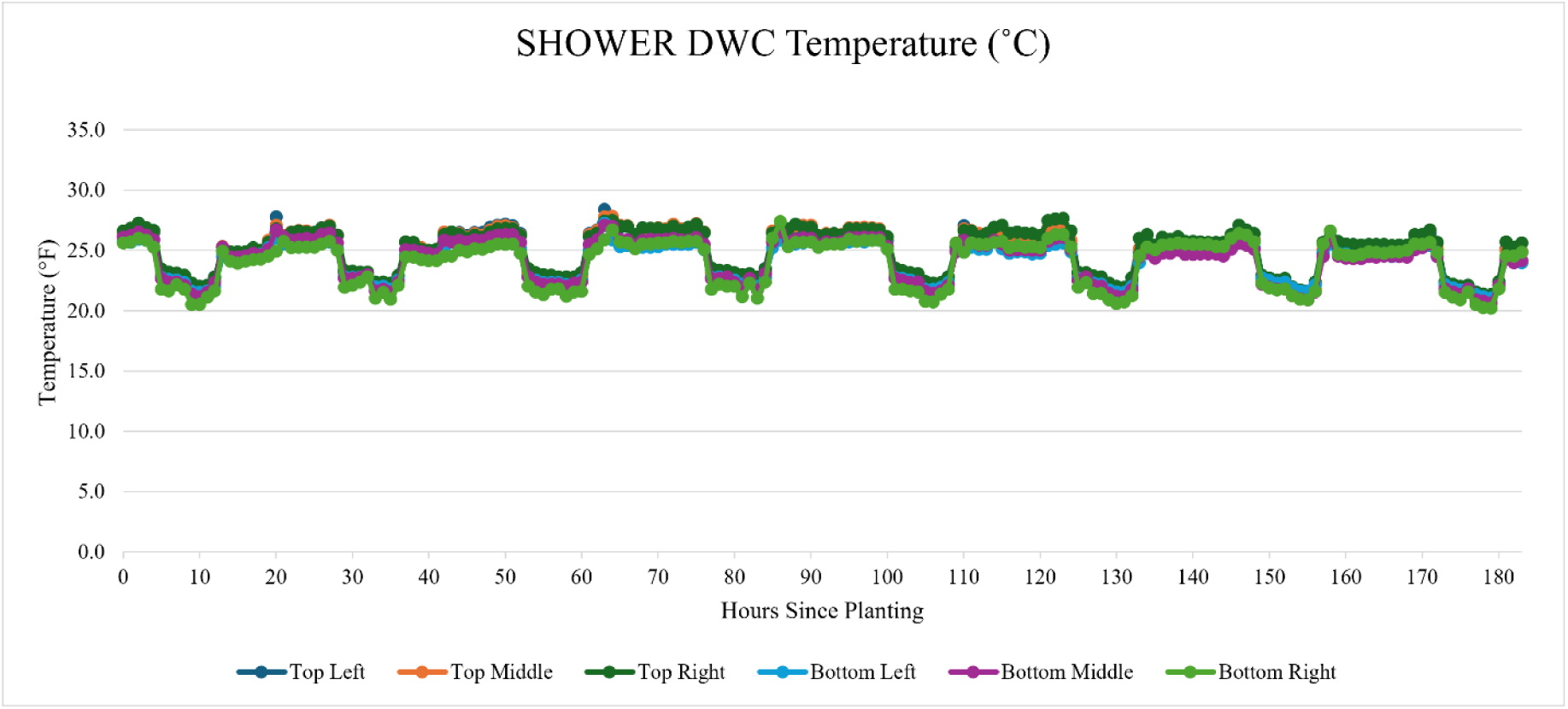
Temperature of the DWC growth chamber for approximately 7 days during the summer months.

**Table 4.** Summarized map of the average environmental conditions experienced by the crops on each growth level.

|  |  |  |
| --- | --- | --- |
| Day/Night Temp (°C):<br>$25.4 \pm 3.1/21.3 \pm 0.6$<br>Day/Night Humidity (%):<br>$63.99 \pm 7.3/80.26 \pm 5.4$<br>PPFD (15.24 cm @ 100%):<br>188.29<br>Wind Speed (m/s): 0.12 | Day/Night Temp (°C):<br>$28.5 \pm 1.5/21.4 \pm 0.8$<br>Day/Night Humidity (%):<br>$63.48 \pm 5.0/78.52 \pm 2.8$<br>PPFD (15.24 cm @ 100%):<br>181.25<br>Wind Speed (m/s): 0.21 | Day/Night Temp (°C):<br>$28.5 \pm 1.4/21.0 \pm 0.5$<br>Day/Night Humidity (%):<br>$60.80 \pm 3.0/82.60 \pm 4.9$<br>PPFD (15.24 cm @ 100%):<br>179.30<br>Wind Speed (m/s): 0.11 |
| Day/Night Temp (°C):<br>$26.9 \pm 1.13/21.7 \pm 0.6$<br>Day/Night Humidity (%):<br>$61.37 \pm 4.8/79.88 \pm 6.0$<br>PPFD (15.24 cm @ 100%):<br>193.76<br>Wind Speed (m/s): 0.22 | Day/Night Temp (°C):<br>$25.7 \pm 2.8/21.3 \pm 0.7$<br>Day/Night Humidity (%):<br>$65.85 \pm 6.1/79.66 \pm 5.4$<br>PPFD (15.24 cm @ 100%):<br>191.59<br>Wind Speed (m/s): 0.19 | Day/Night Temp (°C):<br>$25.6 \pm 2.1/21.3 \pm 0.6$<br>Day/Night Humidity (%):<br>$66.78 \pm 6.4/81.82 \pm 5.2$<br>PPFD (15.24 cm @ 100%):<br>189.76<br>Wind Speed (m/s): 0.16 |
| Day/Night Temp (°C):<br>$27.8 \pm 1.0/21.6 \pm 0.7$<br>Day/Night Humidity (%):<br>$58.93 \pm 4.0/77.82 \pm 3.2$<br>PPFD (15.24 cm @ 100%):<br>188.90<br>Wind Speed (m/s): 0.10 | Day/Night Temp (°C):<br>$28.5 \pm 1.9/21.3 \pm 0.8$<br>Day/Night Humidity (%):<br>$65.74 \pm 4.8/78.68 \pm 2.4$<br>PPFD (15.24 cm @ 100%):<br>182.02<br>Wind Speed (m/s): 0.26 | Day/Night Temp (°C):<br>$24.9 \pm 2.2/21.3 \pm 0.6$<br>Day/Night Humidity (%):<br>$63.46 \pm 5.7/71.94 \pm 1.7$<br>PPFD (15.24 cm @ 100%):<br>192.03<br>Wind Speed (m/s): 0.17 |

### Deep Water Culture (DWC) Growth Chamber

The DWC growth chamber has been utilized for three experiments, two of which have grown germinated seeds up to the point of transplant into the full DWC system. Six Kestrel Drop D2 temperature and humidity loggers were installed, one in the center of each level, with a data collected once per hour. Similar to the microgreen and germination racks, improvements were made after each experiment. Additional sheets of reflective wrapping were installed in various locations to improve the lighting uniformity, an additional oscillating fan was added as the airflow was not adequate, and the exhaust fan speed was adjusted depending on the ambient environment outside the chamber. Depending on experiments during the winter versus summer, the temperature and relative humidity environmental parameters differed due to being reliant on the building’s overall temperature setpoint. In the summer, when the building’s air conditioning was running, the temperature and relative humidity inside the growth chamber were noticeably higher compared to the fall and winter conditions. Figures 9 and 10 display the temperature and humidity, respectively. During the daytime, the temperature and humidity averaged 25.68 °C and 61.84% while during the nighttime the averages were 22.22 °C and 72.15%, shown in Table 5 below.

**Figure 10.**
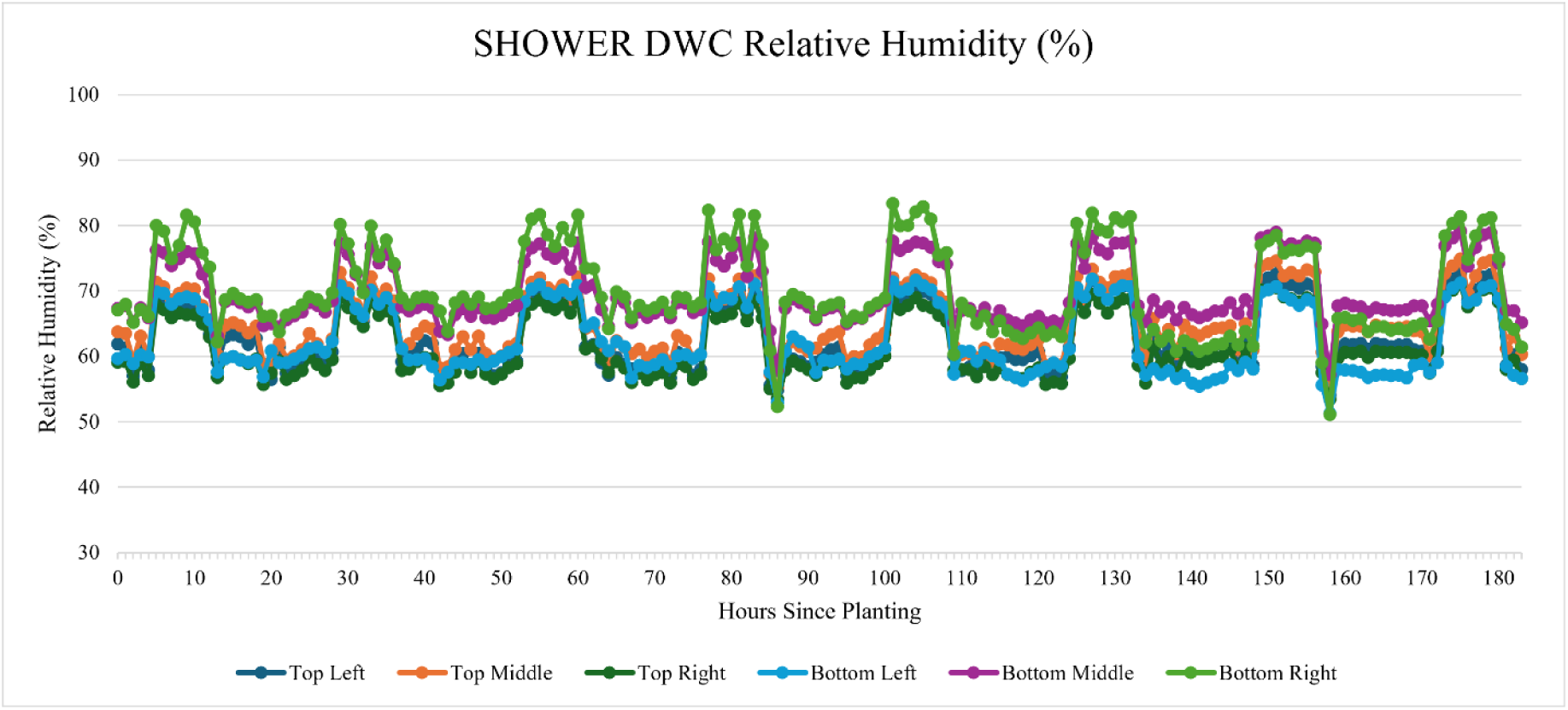
Relative humidity of the DWC growth chamber for approximately 7 days during the summer months.

**Table 5.** Average temperature and relative humidity for the day and night cycles in 24 hours.

|  | Avg Temperature (°C) | StDev | Avg Humidity (%) | StDev |
| --- | --- | --- | --- | --- |
| **Day (11am - 2am)** | 25.70 | 0.81 | 61.84 | 3.97 |
| **Night (3am - 10am)** | 22.22 | 0.69 | 72.15 | 4.30 |

**Table 6.** Light intensity measurements (36) at 100% intensity for the top middle growth level at the height of a plug tray, measured in PPFD.

| Top - Middle - 100% Intensity at Plug Tray Level (PPFD) |  |  |  |  |  |
| --- | --- | --- | --- | --- | --- |
| 207.94 | 228.46 | 200.85 | 205.47 | 226.95 | 202.34 |
| 220.21 | 225.86 | 215.57 | 214.30 | 217.48 | 217.55 |
| 241.10 | 241.32 | 237.31 | 234.16 | 229.08 | 234.41 |
| 240.40 | 240.87 | 235.70 | 232.02 | 228.99 | 235.39 |
| 221.77 | 227.31 | 218.05 | 211.84 | 215.89 | 212.31 |
| 195.10 | 229.21 | 204.48 | 200.76 | 220.11 | 187.58 |
| **Average: 221.06 ± 14.18** |  |  |  |  |  |

The light intensity for the DWC system is measured in PPFD with 36 measurements taken per level, at three separate height intervals, and four distinct levels of intensity. These height intervals include at the top of the plug tray when seeds are germinating and growing in the seedling phase (1.80 in or 4.57 cm), the midway portion between the plug tray and the max canopy height (10.65 in or 27.05 cm), and the max canopy height (21.30 in or 54.10 cm). At each height, measurements were taken at 25%, 50%, 75%, and 100% light intensity. The quantity of measurements ensures a quality overview of each level’s capacity and the intensities between those heights can be interpolated if adjustment is necessary. At 100% intensity, growth of ornamental plant species is optimal even when starting from seed, as 200 PPFD is roughly the desired level for optimal seedling growth and beyond (Degni et al., 2020). Two examples are shown in Tables 7 and 8 below at 100% light intensity, with the four remaining 100% intensity levels shown in S.Tables 4-7.

**Table 7.** Light intensity measurements (36) at 100% intensity for the bottom middle growth level at the height of a plug tray, measured in PPFD.

| Bottom - Middle - 100% Intensity at Plug Tray Level (PPFD) |  |  |  |  |  |
| --- | --- | --- | --- | --- | --- |
| 204.65 | 227.85 | 220.05 | 224.68 | 239.52 | 215.08 |
| 218.33 | 230.48 | 237.36 | 238.10 | 240.14 | 233.84 |
| 221.33 | 228.37 | 235.50 | 236.12 | 238.13 | 236.57 |
| 219.72 | 229.91 | 236.98 | 240.69 | 238.73 | 240.35 |
| 205.62 | 218.82 | 221.12 | 221.70 | 219.84 | 220.44 |
| 182.58 | 212.14 | 215.45 | 210.90 | 203.47 | 205.28 |
| **Average: 224.16 ± 13.59** |  |  |  |  |  |

**Table 8.** DWC growth chamber’s wind speed averaged for each rack, measured in m/s.

| DWC Wind Speed – Averaged (m/s) |  |  |
| --- | --- | --- |
|  | 0.34 |  |
| 0.41 |  | 0.34 |

The wind speed was measured daily between each of the plug trays, at plug tray height, and averaged for the given level. Table 12 displays the wind speed averaged for the entire chamber, and cumulatively was 0.36 ± 0.07 m/s, safely above the 0.2 m/s minimum for proper airflow and mixing.

Figures 11 and 12 below are visual representations of plant growth within the growth chamber during an experiment.

**Figure 11.**
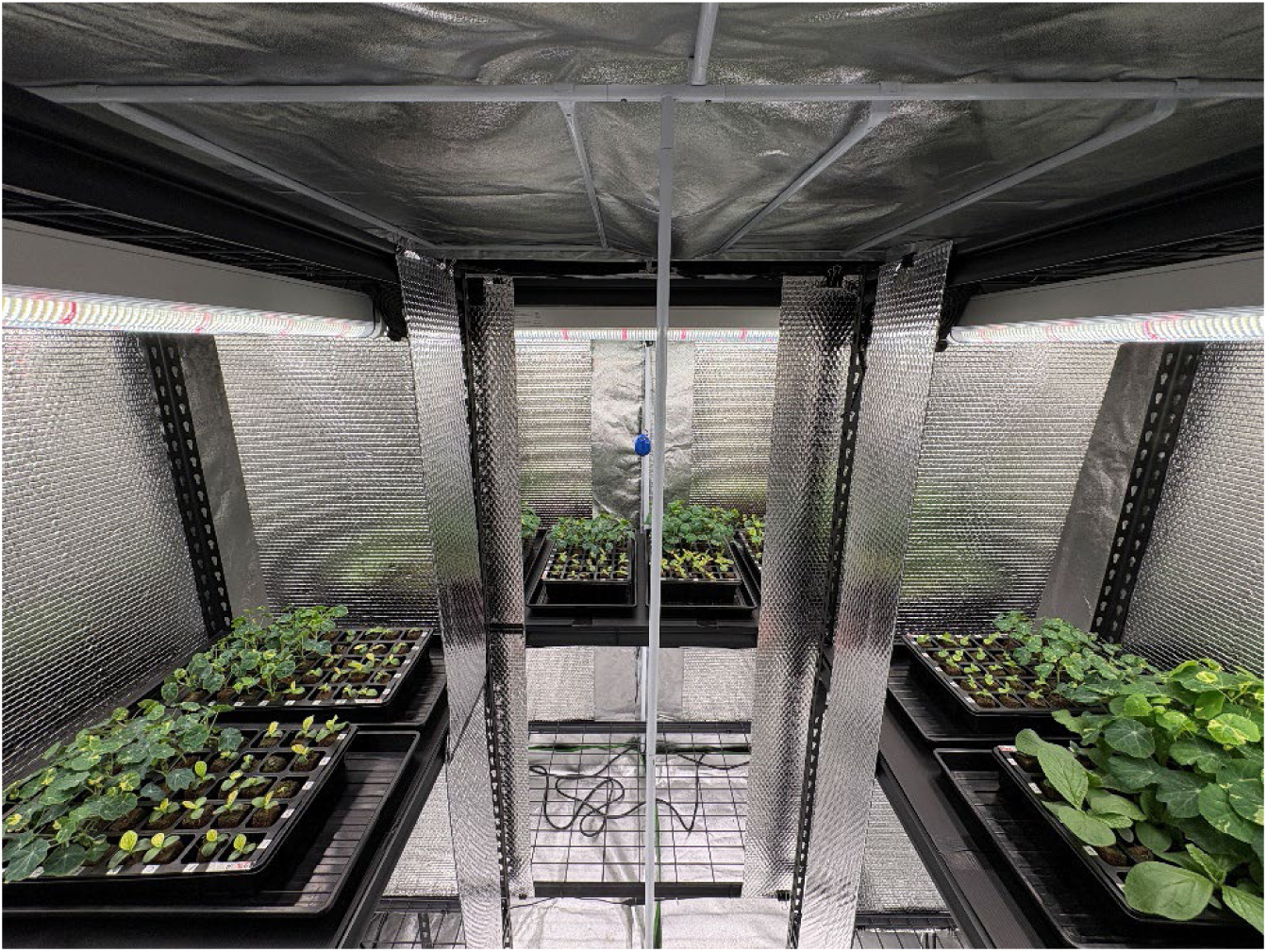
Seedling growth in the DWC growth chamber prior to transplant for full growth.

**Figure 12.**
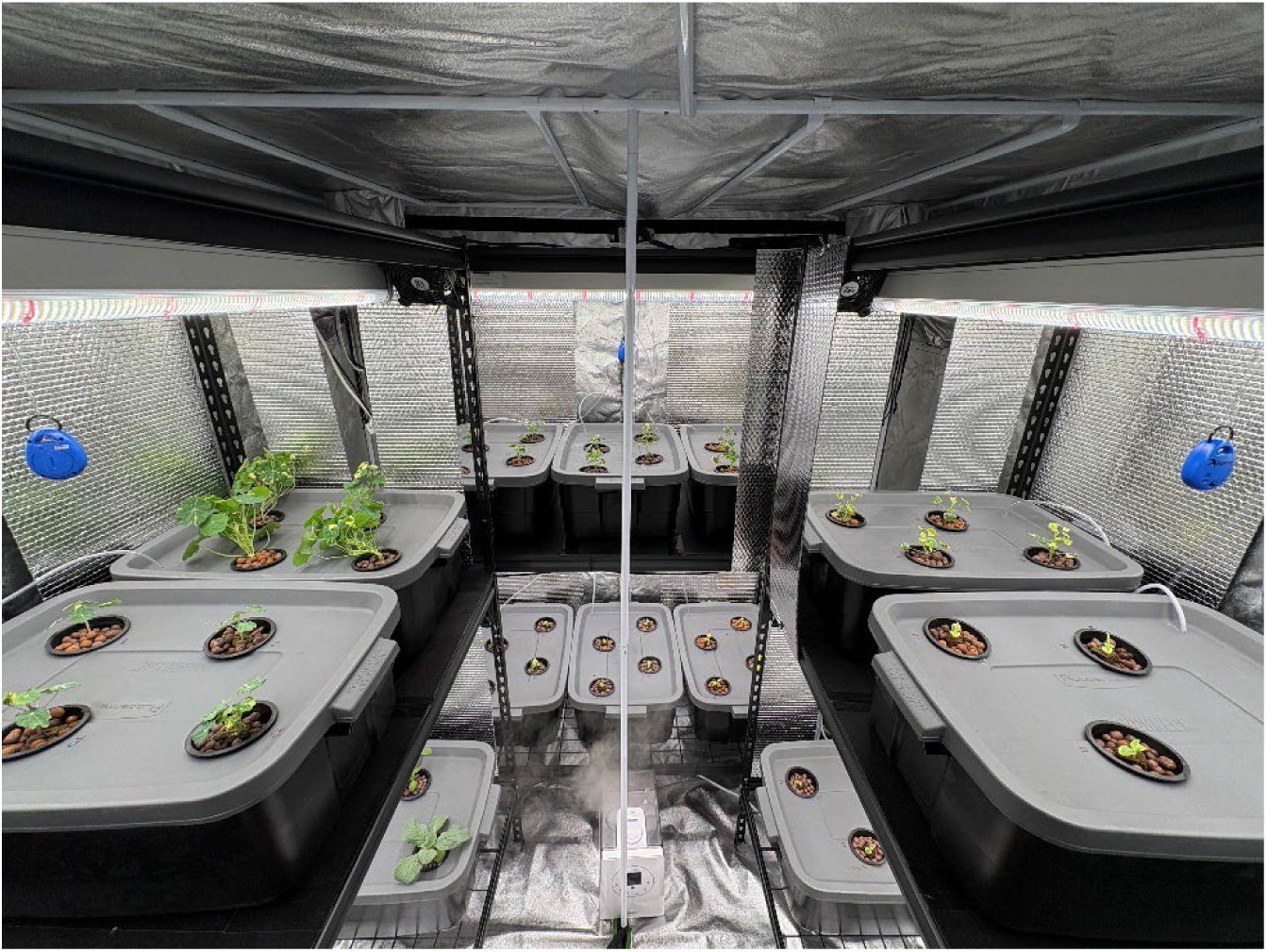
Full growth in the DWC growth chamber after transplant.

### Nutrient Film Technique Grow Tent

The NFT grow tent was tested briefly, for around 1 week, to ensure flow rates and the water levels within the troughs were adequate to support plant growth, as well as to observe any minor differences in environmental conditions on each of the 6 growth levels, before a full 3 weeks of post-transplant growth comprised the NFT data collected. This is more important in the NFT setup, as the individual replicates cannot be spatially rotated through the growth tent, as the microgreen replicates are in the germination and microgreen growth chamber. The Kestrel Datalogger temperature and relative humidity hourly data from the latest experiment are plotted below in Figures 13 and 14. There is one Kestrel datalogger per growth level. The temperature and relative humidity after transplanting the replicates from the Germination/Growth chamber averaged 22.13 ± 0.90 °C and 65 ± 7.1% during the lights-on period and 19.82 ± 0.76 °C and 71 ± 7.1% during the lights-off period.

**Figure 13:**
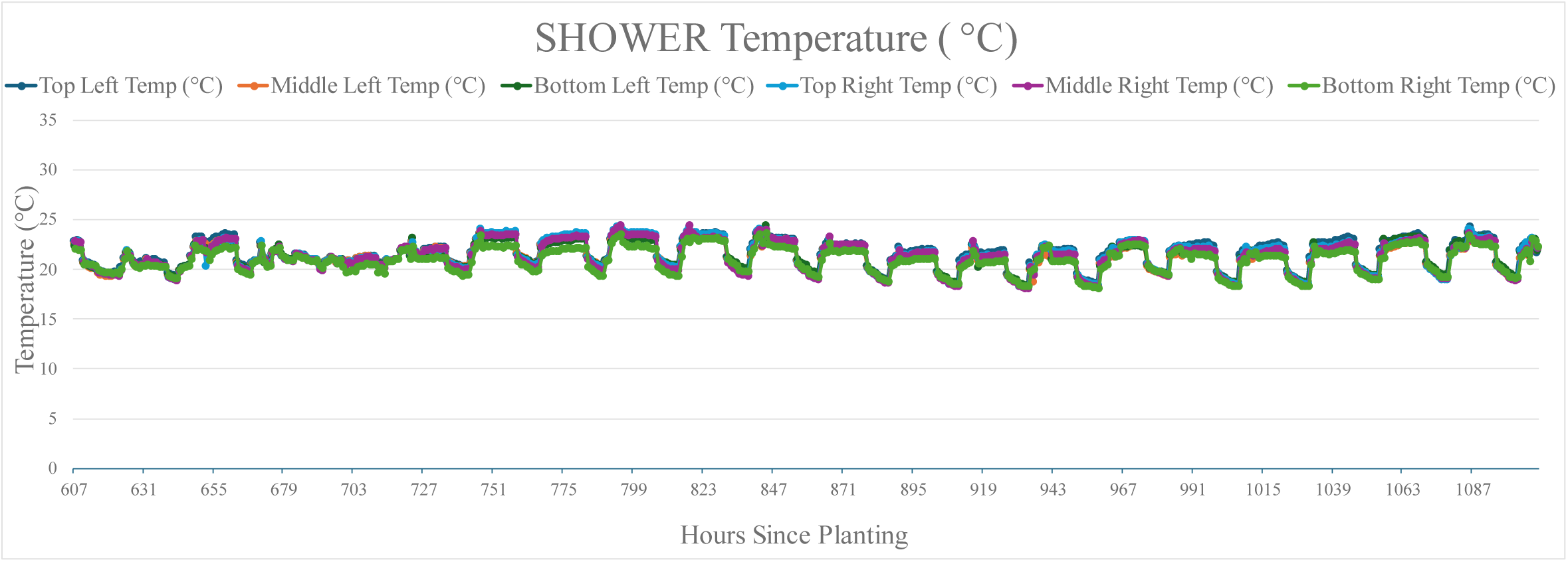
Kestrel temperature data from the NFT experiment.

**Figure 14:**
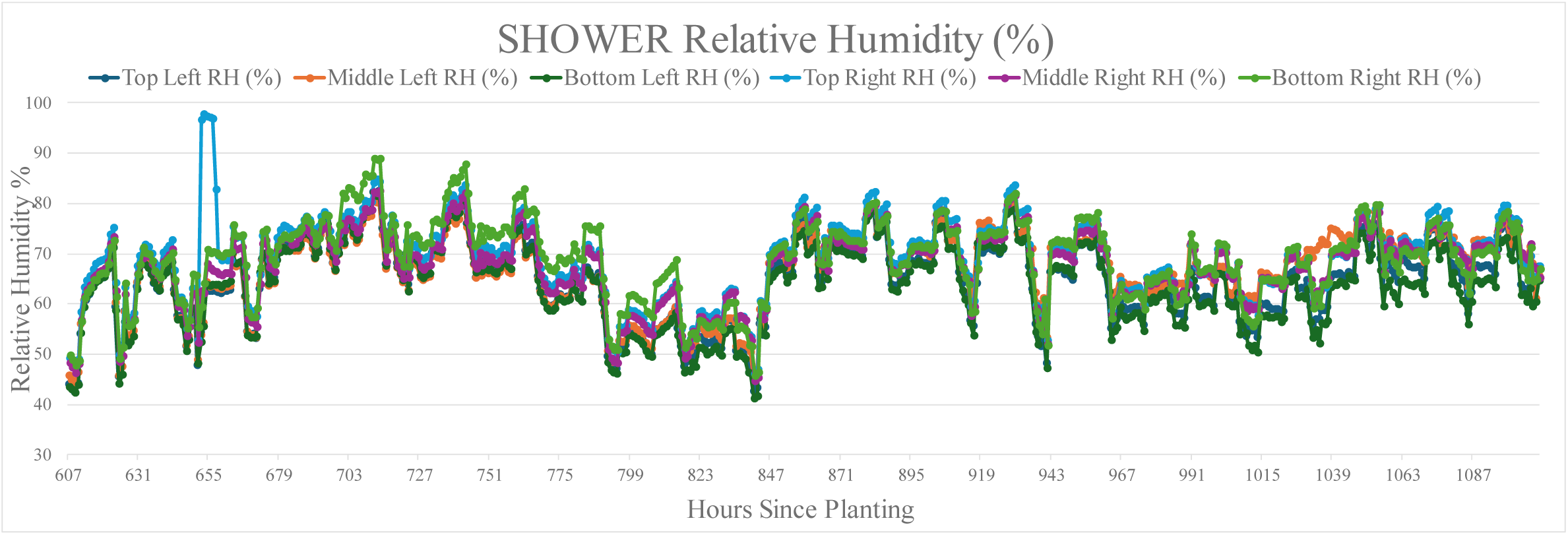
Kestrel relative humidity data from the NFT experiment.

The light intensity (PPFD) was mapped in the NFT grow tent according to the following arrangement: 5 measurements were taken along each 1.22 m PVC trough, corresponding to the plug slots where each replicate would be. There are three troughs per growing level, so there are 15 measurements for a given height and light setting per growth level. Measurements were taken at 50%, 75%, and 100% light intensity settings at heights of trough level, 12.54 cm above the trough, and 30.48 cm above the trough (or just below the LED light bar). The relevant light intensity map for NFT production, 100% intensity at 12.54 cm above the trough level (what the transplanted plugs are exposed to), is provided in supplementary Table 8, but the average PPFD over the whole NFT growth tent is 211.5 ± 11.84 for 12.54 cm above the trough. These light intensity levels are appropriate for leafy green and herb growth, especially as the plants grow towards the light, increasing the intensity they receive, but higher levels are still recommended for flowering/fruiting crop growth, giving reason for the higher light intensity in the DWC growth tent.

The wind speed was measured once a week for each growth level in the center of the middle trough at the average crop canopy height throughout the NFT growth experiment. This was done to ensure measurements were as close to what the plants are experiencing; however, this may have skewed the results slightly, as some irrigation treatments grew better than others, affecting the height of measurements for a given level. The overall NFT growth chamber average wind speed was 0.299 ± 0.11 m/s, and the wind map is shown in supplementary Table 9. This average wind speed is sufficiently above the minimum of 0.2 m/s necessary for uniform air mixing, especially around the crop canopy, facilitating gas exchange and evapotranspiration, enabling the photosynthesis that powers crop growth. All the environmental conditions per growth level within the NFT growth chamber are averaged and presented in Table 9 below.

**Table 9.** Summarized map of the average environmental conditions experienced by the crops at each growth level.

|  |  |  |
| --- | --- | --- |
| Day/Night Temp (°C):<br>$22.5 \pm 0.9/20.0 \pm 0.8$<br>Day/Night Humidity (%):<br>$62.70 \pm 7.28/69.24 \pm 7.54$<br>PPFD (15.24 cm @ 100%): 212.75<br>Wind Speed (m/s): 0.33 | | Day/Night Temp (°C):<br>$22.3 \pm 1.0/19.8 \pm 0.8$<br>Day/Night Humidity (%):<br>$65.82 \pm 7.88/73.34 \pm 6.89$<br>PPFD (15.24 cm @ 100%): 210.26<br>Wind Speed (m/s): 0.30 |
| Day/Night Temp (°C):<br>$22.1 \pm 0.9/19.9 \pm 0.8$<br>Day/Night Humidity (%):<br>$65.42 \pm 7.36/69.58 \pm 7.29$<br>PPFD (15.24 cm @ 100%): 213.94<br>Wind Speed (m/s): 0.46 | | Day/Night Temp (°C):<br>$22.2 \pm 0.9/19.7 \pm 0.8$<br>Day/Night Humidity (%):<br>$65.82 \pm 6.75/71.86 \pm 6.58$<br>PPFD (15.24 cm @ 100%): 211.95<br>Wind Speed (m/s): 0.25 |
| Day/Night Temp (°C):<br>$22.1 \pm 0.9/20.0 \pm 0.8$<br>Day/Night Humidity (%):<br>$61.83 \pm 7.59/68.39 \pm 7.59$<br>PPFD (15.24 cm @ 100%): 209.56<br>Wind Speed (m/s): 0.21 | | Day/Night Temp (°C):<br>$21.7 \pm 0.8/19.6 \pm 0.7$<br>Day/Night Humidity (%):<br>$67.24 \pm 7.43/73.42 \pm 7.22$<br>PPFD (15.24 cm @ 100%): 210.71<br>Wind Speed (m/s): 0.24 |

Figure 15 depicts the NFT growth system in use during the experiment, with the front door unzipped and propped open to see the layout of the experimental treatments.

**Figure 15.**
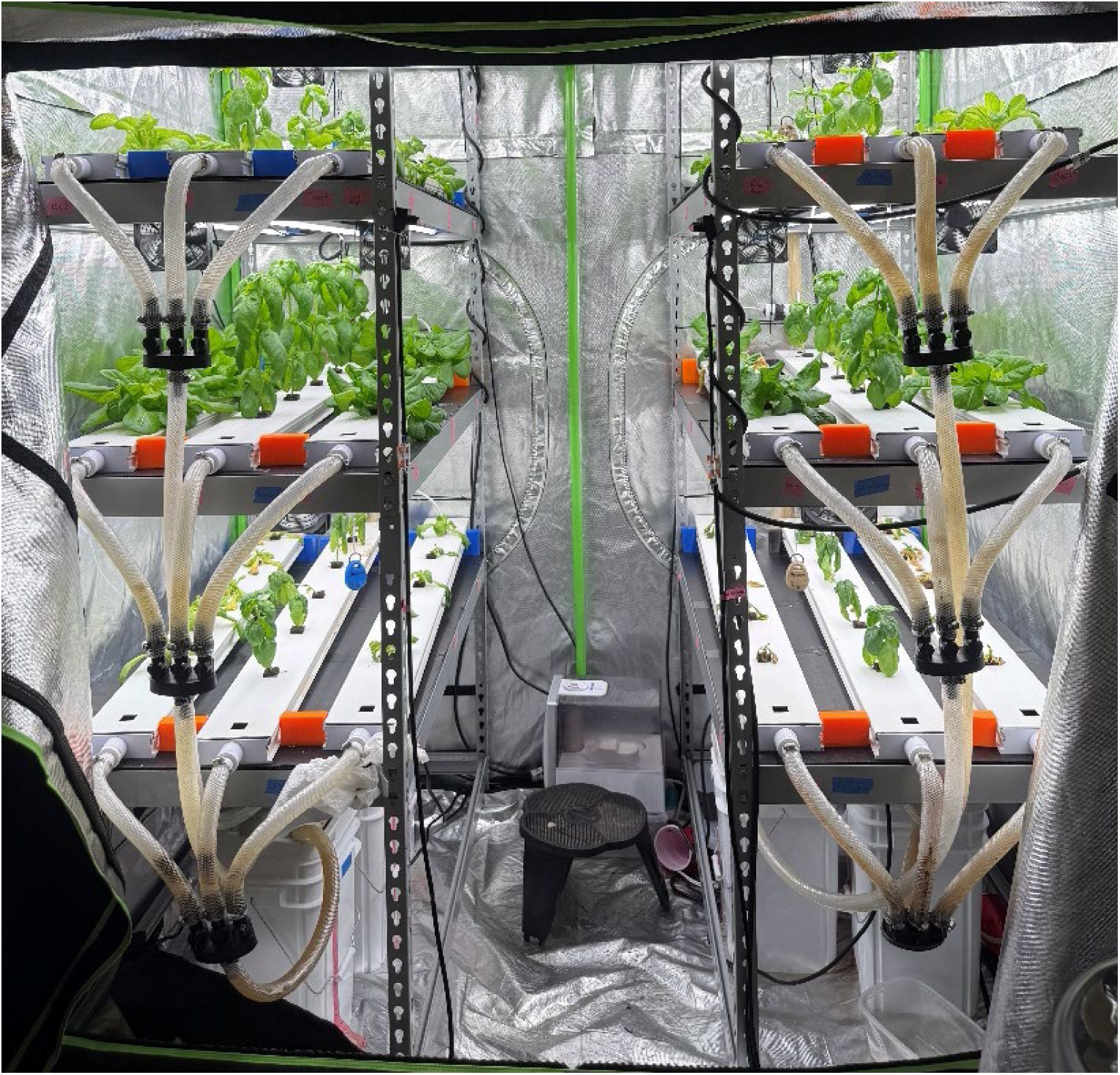
Image of the NFT growth system in use with the front tent flaps open for visibility.

## 4. Discussion

These three growth systems were developed as a lower-cost solution to hydroponic growth chambers that can facilitate research on multiple irrigation solutions at once, in a compact footprint. There are very few environmental control levers that can be pulled except for adjusting the humidifier settings and the circulation and exhaust fans. Despite this, the growth systems maintained consistent performance across different growth levels and across experiments due to being located inside a building that maintains general temperature and humidity control. The designs of these systems also function as a baseline that other users can build from and improve to suit their individual needs, as not everyone is interested in alternative irrigation solutions. A few possible improvements, depending on the research area of interest, include active environmental control feedback with Arduinos, installing imaging/sensor equipment for continuous plant monitoring, adjusting the LED lights for spectrum experiments, and implementing pH and EC dosing systems to continuously provide ideal crop nutrition. The authors hope that other academic researchers, start-up companies, and at-home growers who want to get involved with hydroponic systems but can’t afford the costs associated with high-tech growth chambers or multi-acre greenhouses will leverage the microgreen, DWC, and NFT system designs and parts lists to get started.

## 5. Conclusion

This work can serve as a blueprint or framework for building cost-effective hydroponic systems for both commercial and academic uses. The customization of these systems allows for the production of a wide range of vegetative and ornamental crops using both traditional and nontraditional source waters. The work should serve as a foundation for creating a system that can be continuously improved and modified to the exact needs of the end user. Future improvements may focus on the refilling of humidifier water supplies, automatic crop watering systems, providing a more consistent wind speed over all the plant growth levels within a growth system, and active cooling. These systems consist primarily of off-the-shelf components, with the major items totaling under $2,700 for the microgreens and germination system, $3,700 for the DWC system, and $4,200 for the NFT system. These systems lessen the barriers to entry, enabling many stakeholders to enter the research and production spaces of CEA and better propel the industry forward.

## Abbreviations

CEA: Controlled Environment Agriculture
UA: Urban Agriculture
VF: Vertical Farming
NFT: Nutrient Film Tecnihnique
DWC: Deep Water Culture
RH: Relative Humidity
LED: Light Emitting Diode
PPFD: photosynthetic photon flux density
R&D: Research and Development

## Notes

### Competing Interest Statement

The authors have declared no competing interest.

### Funding

- United States Department of Agriculture: 2023-79000-38974

