## Supplementary Files and Tables for "Design and Construction of Three Customizable Pilot-Scale Hydroponic Systems"

### Supplementary Information

1. .STL file for the side splash guards

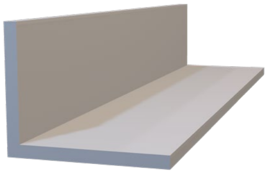

2. .STL file for the outlet splash guards

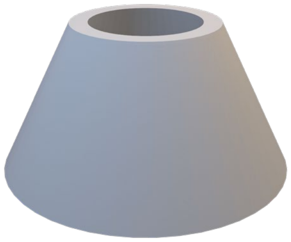

3. .STL file for the outflow side spacers

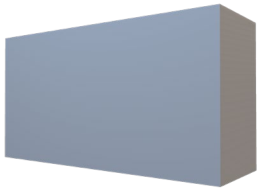

4. .STL file for the inlet side riser-spacers

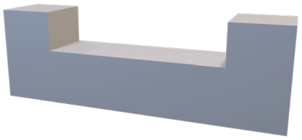

**S.Table 1.** Light map of microgreen growth chamber at 100% intensity at tray level.

| | | | | | | | | |
|---|---|---|---|---|---|---|---|---|
| 145.89 | 155.33 | 153.06 | 150.31 | 147.59 | 153.67 | 152.29 | 153.90 | 150.44 |
| 160.28 | 163.06 | 166.03 | 160.15 | 159.41 | 163.31 | 161.30 | 159.20 | 155.88 |
| 158.47 | 160.47 | 163.74 | 160.93 | 162.88 | 163.24 | 165.83 | 165.39 | 163.36 |
| 149.49 | 155.88 | 157.10 | 161.57 | 159.85 | 159.93 | 153.95 | 154.49 | 154.02 |
| 163.10 | 159.63 | 148.32 | 153.35 | 136.15 | 146.21 | 138.06 | 149.22 | 160.93 |
| 171.11 | 167.06 | 163.67 | 169.88 | 156.69 | 162.21 | 156.73 | 162.29 | 169.32 |
| 172.11 | 168.69 | 167.72 | 174.67 | 164.81 | 167.13 | 166.63 | 167.67 | 173.82 |
| 165.19 | 164.06 | 166.06 | 166.43 | 160.32 | 167.06 | 160.09 | 165.75 | 167.68 |
| 161.76 | 154.94 | 152.29 | 148.93 | 144.72 | 158.40 | 148.50 | 150.56 | 168.5387 |
| 173.23 | 160.40 | 158.45 | 155.22 | 156.08 | 159.42 | 156.81 | 160.86 | 179.6712 |
| 174.39 | 167.22 | 163.77 | 160.46 | 158.50 | 162.04 | 162.50 | 167.76 | 181.4704 |
| 166.32 | 161.39 | 161.48 | 151.77 | 144.22 | 152.67 | 154.91 | 160.61 | 176.7908 |

**S.Table 2.** Light map of the microgreen growth chamber at 100% intensity, 15.24 cm above tray level.

| | | | | | | | | |
|---|---|---|---|---|---|---|---|---|
| 167.16 | 173.74 | 185.59 | 171.60 | 153.33 | 169.38 | 172.74 | 170.72 | 171.37 |
| 194.03 | 178.65 | 183.21 | 195.71 | 185.15 | 193.88 | 184.27 | 172.74 | 187.52 |
| 179.76 | 175.68 | 183.10 | 165.43 | 163.65 | 173.72 | 173.06 | 170.75 | 173.58 |
| 200.09 | 189.16 | 189.32 | 203.41 | 194.92 | 204.86 | 190.32 | 189.68 | 194.86 |
| 198.48 | 197.22 | 203.80 | 185.35 | 177.42 | 201.66 | 199.52 | 180.88 | 188.48 |
| 186.69 | 178.02 | 191.49 | 200.64 | 178.77 | 194.36 | 189.15 | 170.40 | 188.16 |
| 191.40 | 179.83 | 191.29 | 198.54 | 189.19 | 203.23 | 191.28 | 175.11 | 189.58 |
| 195.48 | 196.64 | 214.75 | 204.97 | 173.56 | 191.41 | 207.89 | 191.38 | 205.34 |
| 202.93 | 182.08 | 197.30 | 172.14 | 164.98 | 187.35 | 199.41 | 190.69 | 203.85 |
| 195.00 | 175.64 | 176.38 | 189.80 | 172.18 | 188.61 | 183.57 | 179.20 | 198.96 |
| 188.58 | 172.25 | 183.42 | 176.25 | 169.88 | 184.18 | 180.54 | 174.59 | 186.70 |
| 199.02 | 193.85 | 200.30 | 195.89 | 185.09 | 197.94 | 202.77 | 196.51 | 207.60 |

**S.Table 3.** Heat map of wind speed conditions in the microgreen growth chamber (the circulation fans are situated between the 0.27 and 0.25 cells, along with the 0.47 and 0.29 cells, respectively)

| | | | | | | | | |
|---|---|---|---|---|---|---|---|---|
| 0.16 | 0.10 | 0.12 | 0.36 | 0.14 | 0.14 | 0.15 | 0.08 | 0.10 |
| 0.20 | 0.19 | 0.27 | 0.25 | 0.13 | 0.20 | 0.20 | 0.12 | 0.17 |
| 0.09 | 0.09 | 0.11 | 0.13 | 0.20 | 0.47 | 0.29 | 0.13 | 0.10 |

**S.Table 4.** Light intensity measurements (36) at 100% intensity for the top left growth level at the height of a plug tray, measured in PPFD.

| Top - Left - 100% Intensity at Plug Tray Level (PPFD) | | | | | |
|---|---|---|---|---|---|
| 177.70 | 203.71 | 215.23 | 223.36 | 231.51 | 197.93 |
| 178.70 | 198.30 | 218.49 | 231.25 | 232.97 | 217.14 |
| 183.09 | 205.26 | 228.61 | 239.90 | 242.08 | 232.57 |
| 182.93 | 204.76 | 225.18 | 234.68 | 240.18 | 233.65 |
| 172.09 | 189.41 | 215.28 | 214.27 | 228.83 | 223.46 |
| 162.10 | 172.50 | 190.90 | 198.42 | 216.94 | 227.66 |
| **Average: 210.86 ± 22.48** | | | | | |

**S.Table 5.** Light intensity measurements (36) at 100% intensity for the top right growth level at the height of a plug tray, measured in PPFD.

| Top - Right - 100% Intensity at Plug Tray Level (PPFD) | | | | | |
|---|---|---|---|---|---|
| 190.79 | 227.36 | 224.52 | 223.65 | 201.69 | 178.87 |
| 210.82 | 228.00 | 226.78 | 220.62 | 204.99 | 181.37 |
| 231.19 | 242.55 | 244.03 | 235.30 | 215.78 | 191.12 |
| 233.24 | 241.48 | 235.39 | 226.08 | 205.30 | 185.77 |
| 216.30 | 217.47 | 212.76 | 206.42 | 192.12 | 172.51 |
| 228.58 | 218.00 | 202.56 | 194.25 | 179.68 | 160.31 |
| **Average: 211.32 ± 21.57** | | | | | |

**S.Table 6.** Light intensity measurements (36) at 100% intensity for the bottom left growth level at the height of a plug tray, measured in PPFD.

| Bottom - Left - 100% Intensity at Plug Tray Level (PPFD) | | | | | |
|---|---|---|---|---|---|
| 184.06 | 214.93 | 236.78 | 243.75 | 252.26 | 218.19 |
| 194.11 | 216.34 | 242.35 | 256.74 | 258.22 | 238.59 |
| 196.92 | 224.91 | 242.82 | 249.56 | 250.20 | 235.83 |
| 193.86 | 231.42 | 247.50 | 250.25 | 254.63 | 241.35 |
| 204.23 | 226.24 | 242.30 | 244.02 | 252.31 | 238.45 |
| 200.29 | 220.91 | 235.13 | 242.38 | 242.61 | 239.62 |
| **Average: 232.34 ± 20.06** | | | | | |

**S.Table 7.** Light intensity measurements (36) at 100% intensity for the bottom right growth level at the height of a plug tray, measured in PPFD.

| Bottom - Right - 100% Intensity at Plug Tray Level (PPFD) | | | | | |
|---|---|---|---|---|---|
| 203.05 | 239.70 | 227.73 | 240.52 | 213.08 | 182.14 |
| 220.76 | 254.12 | 246.52 | 236.02 | 224.10 | 192.93 |
| 235.36 | 256.20 | 254.08 | 243.87 | 233.59 | 205.35 |
| 230.44 | 245.36 | 243.55 | 234.25 | 226.86 | 202.56 |
| 228.94 | 238.17 | 241.66 | 234.38 | 226.68 | 200.26 |
| 240.37 | 227.86 | 243.75 | 235.92 | 219.85 | 198.17 |
| **Average: 228.56 ± 18.29** | | | | | |

**S.Table 8.** Light Map of NFT Growth Chamber at 100% intensity, 12.54 cm above Trough Level

| | | |
|---|---|---|
| 194.6 | 214.86 | 203.2 |
| 216.8 | 233.95 | 213.4 |
| 215.7 | 230.83 | 203.5 |
| 213.9 | 232.1 | 194.9 |
| 212.4 | 225.2 | 185.9 |
| 197.2 | 207.35 | 190.2 |
| 217.8 | 232.26 | 220.7 |
| 212.9 | 228.13 | 219 |
| 207.7 | 227.6 | 215.7 |
| 200 | 218.56 | 213.9 |
| 185.7 | 209.97 | 196.4 |
| 198.2 | 221.62 | 229.6 |
| 199.4 | 217.23 | 218.9 |
| 204.3 | 219.86 | 214.6 |
| 209 | 214.18 | 204.5 |

| | | |
|---|---|---|
| 201.3 | 217.1 | 207 |
| 209.1 | 231.4 | 213.6 |
| 197 | 222.9 | 208.4 |
| 193.3 | 224 | 212.4 |
| 185 | 219.4 | 212.1 |
| 186.4 | 215.1 | 200.3 |
| 211.8 | 232.1 | 222.7 |
| 208.6 | 223.8 | 213.6 |
| 206.1 | 223.1 | 212.6 |
| 207 | 216 | 200 |
| 186.3 | 210.1 | 192.7 |
| 220.3 | 226.6 | 214.8 |
| 215.9 | 222.3 | 208.3 |
| 211.3 | 219 | 207.9 |
| 202.8 | 214 | 208 |

**S.Table 9.** Heat map of wind speed conditions in the NFT Growth Chamber

| |
|---|
| 0.334 |
| 0.463 |
| 0.205 |

| |
|---|
| 0.304 |
| 0.249 |
| 0.238 |
